# Dynamics of allosteric regulation of the phospholipase C-γ isozymes upon recruitment to membranes

**DOI:** 10.1101/2022.02.23.481568

**Authors:** Edhriz Siraliev-Perez, Jordan T.B. Stariha, Reece M. Hoffmann, Brenda R. Temple, Qisheng Zhang, Nicole Hajicek, Meredith L Jenkins, John E. Burke, John Sondek

## Abstract

Numerous receptor tyrosine kinases and immune receptors activate phospholipase C-gamma (PLC-γ) isozymes at membranes to control diverse cellular processes including phagocytosis, migration, proliferation, and differentiation. The molecular details of this process are not well understood. Using hydrogen-deuterium exchange mass spectrometry (HDX-MS), we show that PLC-γ1 is relatively inert to lipid vesicles that contain its substrate, PIP_2_, unless first bound to the kinase domain of the fibroblast growth factor receptor (FGFR1). Exchange occurs throughout PLC-γ1 and is exaggerated in PLC-γ1 containing an oncogenic substitution (D1165H) that allosterically activates the lipase. These data support a model whereby initial complex formation shifts the conformational equilibrium of PLC-γ1 to favor activation. This receptor-induced priming of PLC-γ1 also explains the capacity of a kinase-inactive fragment of FGFR1 to modestly enhance the lipase activity of PLC-γ1 operating on lipid vesicles but not a soluble analog of PIP_2_ and highlights cooperativity between receptor engagement and membrane proximity. Priming is expected to be greatly enhanced for receptors embedded in membranes and nearly universal for the myriad of receptors and co-receptors that bind the PLC-γ isozymes.

## Introduction

The 13 mammalian phospholipase C (PLC) isozymes hydrolyze the minor membrane phospholipid phosphatidylinositol 4,5-bisphosphate (PIP_2_) to create the second messengers inositol 1,4,5-trisphosphate (IP_3_) and diacylglycerol (DAG) (Gresset et al., 2012; Kadamur and Ross, 2013). IP_3_ diffuses throughout the cytosol and binds to IP_3_-gated receptors in the endoplasmic reticulum, resulting in an increase in the concentration of intracellular calcium ions. In contrast, DAG remains embedded in the inner leaflet of the plasma membrane, where it recruits and activates multiple signaling proteins containing C1 domains including the conventional isoforms of protein kinase C (PKC). In addition, DAG is a precursor of phosphatidic acid, itself an important signaling lipid. PLC-dependent depletion of PIP_2_ levels in the plasma membrane also regulates the activity of numerous ion channels and signaling proteins. Accordingly, the PLCs orchestrate fluctuations in the abundance of both phospholipids and second messengers to control important biological processes including proliferation (Noh et al., 1995) and cell migration (Asokan et al., 2014).

The PLC-γ isozymes, PLC-γ1 and PLC-γ2, are activated downstream of myriad cell surface receptors and regulate signaling cascades that modulate multiple aspects of embryonic development, directed cell migration, and the immune response. Mounting evidence also indicates that the PLC-γ isozymes are drivers of human disease including cancer and immune disorders (Koss et al., 2014). For example, PLC-γ1 is overexpressed and presumably activated in breast cancer (Arteaga et al., 1991). In addition, genome-wide sequencing studies have identified somatic gain-of-function mutations in PLC-γ1 and PLC-γ2 in a variety of immunoproliferative malignancies. In one cohort of patients with adult T cell leukemia/lymphoma, PLC-γ1 is the most frequently mutated gene with almost 40% of patients harboring at least one substitution in the isozyme (Kataoka et al., 2015). Moreover, mutations in PLC-γ2 arise in ~30% of patients with relapsed chronic lymphocytic leukemia after treatment with ibrutinib, a covalent inhibitor of Bruton’s tyrosine kinase (Woyach et al., 2014). Recently, a putative gain-of-function variant was identified in PLC-γ2, Pro522Arg, that was associated with a decreased risk of late-onset Alzheimer’s disease (Sims et al., 2017). This variant also seems to slow cognitive decline in patients with mild cognitive impairment (Kleineidam et al., 2020), suggesting that in some contexts, elevated PLC-γ activity is beneficial.

Among the PLCs, PLC-γ1 and PLC-γ2 are distinct in that they are the only isozymes directly activated by tyrosine phosphorylation. It is widely accepted that phosphorylation of Tyr783 in PLC-γ1 and the equivalent site in PLC-γ2, Tyr759, is required for their regulated activation (Kim et al., 1991; Kim et al., 1990; Watanabe et al., 2001). Phosphorylation and consequently, activation of the PLC-γ isozymes is mediated by two major classes of tyrosine kinases. Multiple receptor tyrosine kinases (RTKs) including receptors for growth factors such as fibroblast growth factor (Burgess et al., 1990), as well as Trk receptors (Vetter et al., 1991) activate the PLC-γ isozymes. PLC-γ1 and PLC-γ2 are also phosphorylated by soluble tyrosine kinases associated with immune receptors, including members of the Src, Syk, and Tec families (Law et al., 1996; Nakanishi et al., 1993; Schaeffer et al., 1999).

PLC-γ1 and PLC-γ2 contain a distinctive array of regulatory domains that controls their phosphorylation-dependent activation. The phosphorylation sites required for lipase activity (Tyr783 in PLC-γ1 and Tyr759 in PLC-γ2) are located within this array (Gresset et al., 2010; Kim et al., 1991; Nishibe et al., 1990). The array also harbors two SH2 domains, nSH2 and cSH2, which mediate binding of tyrosine kinases (Nishibe et al., 1990), and autoinhibition of lipase activity (Gresset et al., 2010), respectively. Indeed, deletion or mutation of the cSH2 domain is sufficient to constitutively activate PLC-γ1 *in vitro* and in cells (Gresset et al., 2010; Hajicek et al., 2013). Importantly, the release of autoinhibition mediated by the cSH2 domain is coupled directly to the phosphorylation of Tyr783 (Gresset et al., 2010). A PLC-γ1 peptide encompassing phosphorylated Tyr783 (pTyr783) binds to the isolated cSH2 domain with high affinity (K_D_ ~360 nM). In the context of the full-length isozyme, mutation of the cSH2 domain such that it cannot engage phosphorylated tyrosines ablates kinase-dependent activation of PLC-γ1 *in vitro*. Presumably, the intramolecular interaction between pTyr783 and the cSH2 domain drives a substantial conformational rearrangement resulting in activation of the enzyme. In addition to these regulatory components, the array also harbors scaffolding properties mediated by a split pleckstrin homology (sPH) domain and an SH3 domain that bind multiple signaling and adaptor proteins including the monomeric GTPase Rac2 (Walliser et al., 2008) and SLP-76 (Yablonski et al., 2001), respectively.

The X-ray crystal structure of essentially full-length PLC-γ1 highlights how the regulatory array integrates these myriad functions (Hajicek et al., 2019) (**Fig. 1A-B**). In the autoinhibited state, the array is positioned directly on top of the catalytic core of the enzyme. This arrangement, coupled with the overall electronegative character of the array effectively blocks the lipase active site from spuriously engaging the plasma membrane. Two major interfaces anchor the regulatory array to the catalytic core. The first interface is formed between the cSH2 domain and the C2 domain, and essentially buries the surface of the cSH2 domain that engages the region encompassing pTyr783. The second interface is comprised of the sPH domain and the catalytic TIM barrel. More specifically, the sPH domain sits atop the hydrophobic ridge in the TIM barrel, effectively capping the ridge and preventing its insertion into membranes, a requisite step in the interfacial activation of PLC isozymes. In contrast, the phosphotyrosine-binding pocket on the nSH2 domain, as well as the polyproline-binding surface on the SH3 domain are solvent exposed, implying that these elements are pre-positioned to efficiently engage activated RTKs and adaptor proteins, respectively. The overall architecture of the regulatory array is consistent with an enzyme that is unable to access membrane-resident substrate in the basal state. Upon activation of a RTK, e.g. fibroblast growth factor receptor 1 (FGFR1), the nSH2 domain mediates recruitment of PLC-γ1 to the phosphorylated tail of the receptor, and PLC-γ1 is subsequently phosphorylated by FGFR1 on Tyr783. pTyr783 then binds to the cSH2 domain, unlatching it from the catalytic core. This binding event is likely coupled to additional structural rearrangements of the regulatory array with respect to the catalytic core that ultimately allow the active site to access PIP_2_ embedded in membranes (Hajicek et al., 2019; Liu et al., 2020).

**Figure 1.**
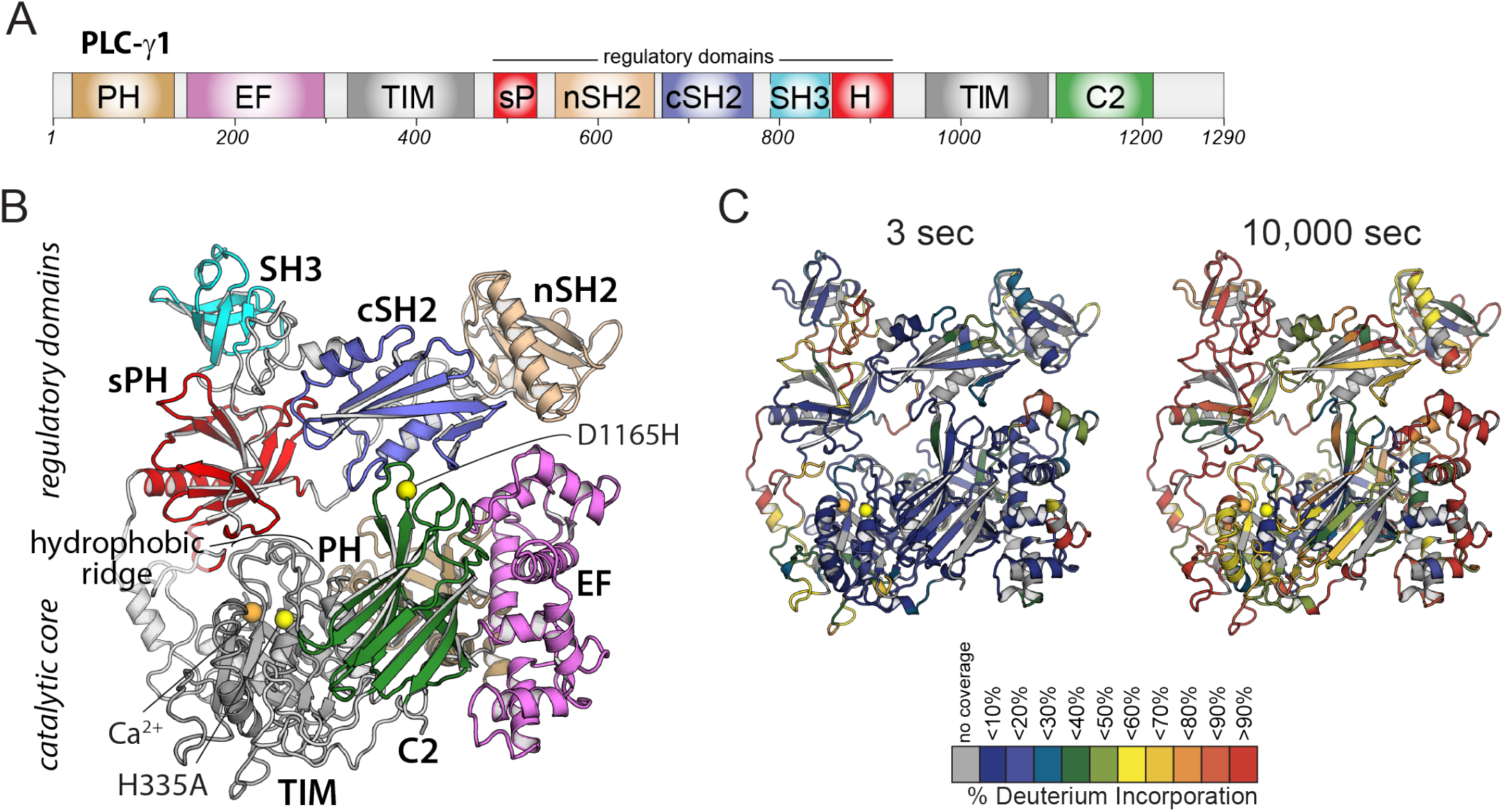
Domain architecture of PLC-γ1 and initial HDX-MS time course. (**A**) Domain schematic of PLC-γ1. Full-length PLC-γ1 possesses a set of regulatory domains inserted within its catalytic core (upper panels). (**B**) Structure of PLC-γ1. In the basally autoinhibited state shown here (PDB ID: 6PBC), the regulatory domains prevent access to membranes by the catalytic core demarked by a Ca^2+^ cofactor (orange). His335 is also within the catalytic core and its substitution (H335A) renders the isozyme catalytically inactive (yellow). Asp1165 substitution (D1165H) disrupts the interface between the core and the regulatory array leading to a constitutively active lipase (also yellow). (**C**) Relative levels of deuterium incorporation are colored on the structure of PLC-γ1 (residue 21 - 1215) according to the legend at either 3 or 10,000 seconds of D_2_O exposure.

Although the core aspects of autoinhibition and phosphorylation-dependent regulation are relatively well established, less is known about the chronology of events subsequent to receptor binding that result in lipase activation. A comparison of the structure of autoinhibited PLC-γ1 and a structure of the tandem SH2 domains of PLC-γ1 bound to the phosphorylated kinase domain of FGFR1 (Bae et al., 2009) suggests a cogent sequence of events. In autoinhibited PLC-γ1, kinase engagement by the nSH2 domain would partially expose the surface of the cSH2 domain that binds pTyr783. Binding of pTyr783 would then fully displace the cSH2 domain from the catalytic core, presumably initiating and propagating the larger structural rearrangements that disrupt the remainder of the autoinhibitory interface, culminating in membrane engagement. Cooperativity between nSH2 and cSH2 domains of PLC-γ1 has been described previously (Bunney et al., 2012), however the functional consequences of this cooperativity were linked to a reduction in the affinity for FGFR1 with no corresponding effect on lipase activity.

Here, we use hydrogen-deuterium exchange mass spectrometry (HDX-MS) to define changes in PLC-γ1 dynamics upon engagement of PIP_2_-containing liposomes and the phosphorylated kinase domain of FGFR1. PLC-γ1 was largely quiescent in the presence of liposomes. In contrast, rapid and widespread changes in exchange kinetics were observed in a PLC-γ1-FGFR1 complex. These changes were centered on the nSH2 domain, and also encompassed large swaths of the autoinhibitory interface. Changes in this exchange profile were further amplified in PLC-γ1 bound to both FGFR1 and liposome. These results are consistent with *in vitro* lipase assays demonstrating that engagement of FGFR1 was sufficient to elevate PLC-γ1 activity, and suggest a model in which receptors and membranes act cooperatively to prime the lipase for full activation.

## Results

### Wide-spread changes in the hydrogen-deuterium exchange of PLC-γ1 by kinase and liposomes

In order to assess the dynamics of PLC-γ1 using HDX-MS, it was first necessary to establish optimized conditions for protein coverage, a suitable time course of H/D exchange, and constructs suitable for membrane binding. These parameters were addressed in two steps. First, optimization of HDX-MS conditions allowed for the generation of 254 peptides covering ~92% of the PLC-γ1 sequence, with an average peptide length of 13 amino acids (**Table S1**). H/D exchange was carried out at five time points (3, 30, 300, 3000, and 10000 sec), which allowed for the interrogation of differences in both highly dynamic and stable regions of secondary structure (**Fig 1C**). Second, it was necessary to design and ensure a system that would allow PLC-γ1 to interact with PIP_2_-containing liposomes over the time course of the exchange reaction without the catalytic conversion of the liposomes. This outcome was accomplished by introduction of a single substitution (H335A) into PLC-γ1 of a catalytic residue necessary for PIP_2_ hydrolysis (**Fig. 1B**). Although several substitutions of active site residues are known to render PLCs catalytically inactive (Ellis et al., 1998), His335 was specifically chosen for its role in the enzymatic process. More specifically, based on analogy to PLC-δ1, His335 is predicted to coordinate the incoming water molecule required for PIP_2_ hydrolysis; it is predicted to be required during the transition state but should have little effect on the affinity of PLC-γ1 for PIP_2_ as a substrate. In addition, His335 does not ligate the calcium ion cofactor in the crystal structure of PLC-γ1 and for this reason is unlikely to perturb the overall structure of the active site.

Consequently, PLC-γ1 (H335A) was purified and its lack of catalytic activity verified using a fluorescent substrate embedded within liposomes (**Fig. S1A**). In addition, a flotation assay was used to show that PLC-γ1 (H335A) and wild-type PLC-γ1 bound PIP_2_-containing liposomes with approximately equal propensity (~20%) (**Fig. S1B**). For flotation measurements, proteins were initially treated to chelate calcium and free calcium concentrations were maintained below 400 nM to prevent substantial PIP_2_ hydrolysis by wild-type PLC-γ1.

Once we determined PLC-γ1 was amenable to studies by HDX-MS, changes in the hydrogen-deuterium exchange profile of PLC-γ1 (H335A) were measured in response to either: i) PIP_2_-containing lipid vesicles, ii) the kinase domain of fibroblast growth factor receptor 1 (FGFR1K) phosphorylated at Tyr766, or iii) both vesicles and phosphorylated kinase (**Fig. 2**). Tyrosine 766 is a major site of phosphorylation in FGFR1 and phosphorylated Tyr766 directly binds PLC-γ1 to recruit it to membranes in cells (Eswarakumar et al., 2005; Mohammadi et al., 1991). Phosphorylated Tyr766 of FGFR1K is also the major site of interaction between the kinase domain and the tandem SH2 array of PLC-γ1 in a crystal structure of these fragments in complex (Bae et al., 2009). Consequently, the kinase domain of FGFR1 was mutated to remove minor sites of tyrosine phosphorylation and to render the kinase resistant to phosphorylation-dependent activation before phosphorylation of Tyr766 and stable, 1:1 complex formation with PLC-γ1 (H335A) (**Figs. S2-S3**).

**Figure 2.**
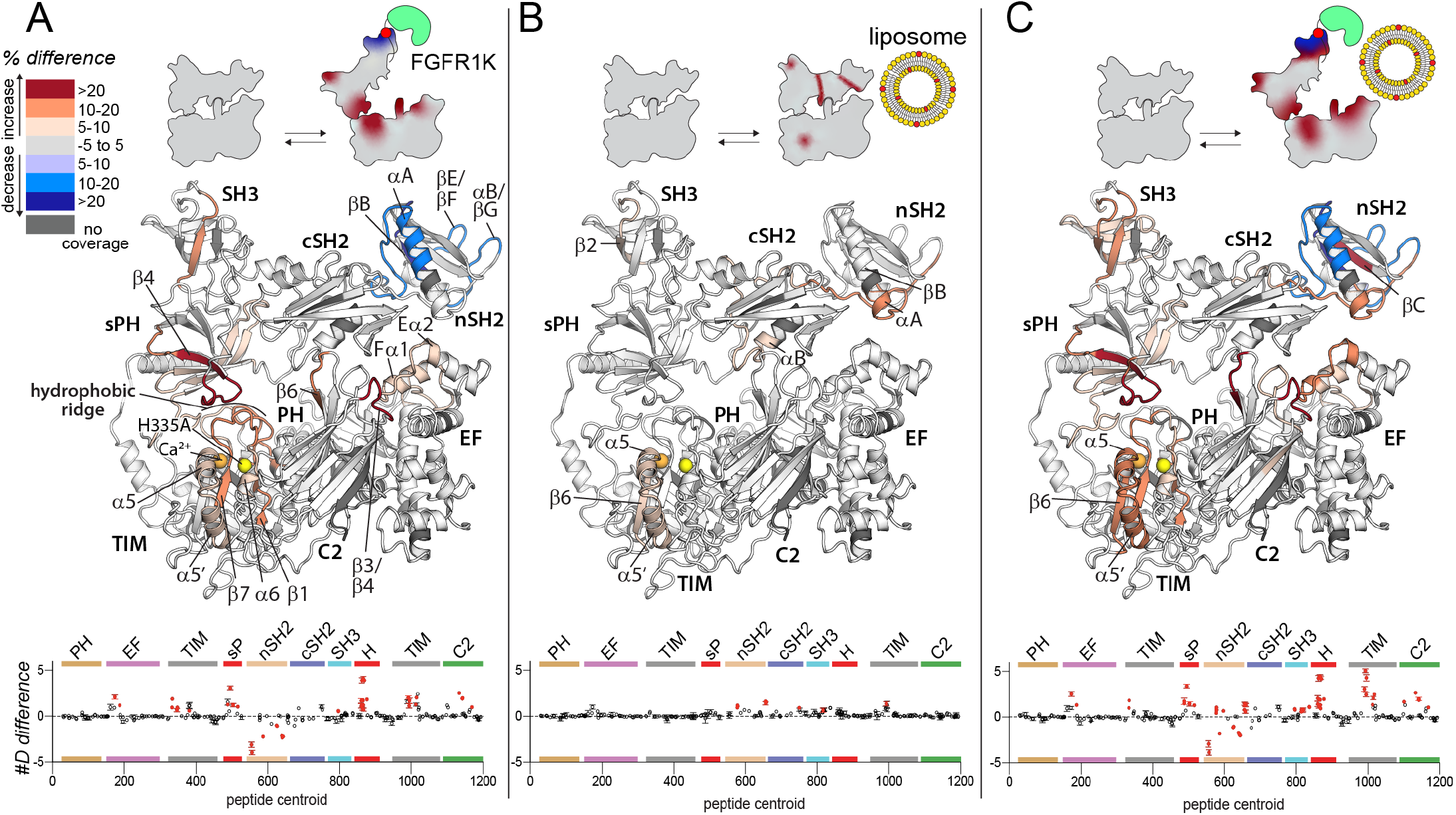
Wide-spread changes in deuterium exchange of PLC-γ1 upon binding FGFR1K. Significant differences in deuterium incorporation are mapped on the structure of PLC-γ1 (H335A) according to the legend for the following three states: bound to phosphorylated FGFR1K (**A**), in the presence of liposomes containing PE:PIP_2_ (90:10) (**B**), or with both kinase and liposomes (**C**). Significant differences in any peptide required three specific conditions: greater than both a 5% and 0.4 Da difference in exchange at any timepoint, and a two-tailed unpaired t-test of p<0.01. The #D difference for each condition is graphed below, which show the total difference in deuterium incorporation over the entire HDX time course, with each point indicating a single peptide (error shown as SD [n=3]). Each circle represents the central residue of a corresponding peptide, with the full deuterium exchange information for all peptides available in the source data. Individual peptides with a significant difference as defined above are colored red.

#### FGFR1K potentially disrupts autoinhibition

There were widespread, significant differences in H/D exchange that occurred in PLC-γ1 (H335A) upon binding phosphorylated FGFR1K (**Fig. 2A**). For all reported significant changes in H/D exchange, three specific conditions had to be met: greater than both a 5% and 0.4 Da difference in exchange at any timepoint, and a two-tailed unpaired t-test of p<0.01. The full HDX data, and differences are reported in the **source data**.

Only the nSH2 domain of PLC-γ1 showed significantly decreased exchange upon binding FGFR1K. It is this domain that directly engages phosphorylated Tyr766 in the crystal structure of the kinase domain of FGFR1 bound to the tandem SH2 domains of PLC-γ1 (**Fig. S4**) (Bae et al., 2009). Consequently, the most straightforward interpretation of the reduced exchange in the complex is that it reflects the major sites of interaction between phosphorylated FGFR1 and PLC-γ1. In fact, regions of reduced exchange within the nSH2 domain match exceptionally well to those regions that directly interact with the phosphorylated portion of FGFR1K in the crystal structure. These regions of the nSH2 domain include the N-terminal portion of the αA helix and the C-terminal portion of the βB strand that interact directly with phosphorylated Tyr766 as well as the βE/βF and αB/βG loops that interact with regions of FGFR1K proximal to phosphorylated Tyr766 (**Figs. 2A, S4-S5**). These data are consistent with previously reported differences in H/D exchange in PLC-γ1 upon binding FGFR1K (Liu et al., 2020).

In contrast to decrease exchange localized to the nSH2 domain, exchange was significantly increased throughout much of PLC-γ1 (H335A) upon binding FGFR1K (**Fig. 2A, S5**). In particular, increased exchange was widespread throughout the interface between the regulatory array and the catalytic core required for autoinhibition of lipase activity. For example, on one side of the interface, all of the loops within the TIM barrel that comprise the hydrophobic ridge showed increased exchange. This increased exchange was mirrored on the other side of the interface by widespread increases in exchange of most of the split PH domain. In fact, the entire first half of the sPH domain along with the first strand (β4) of the second half of the sPH domain showed increased exchange. Indeed, some of the largest increases in percent exchange were found within the β4 strand and the immediately preceding loop that interacts with the hydrophobic ridge of the TIM barrel in the autoinhibited form of PLC-γ1. Loops of the C2 domain interacting with the cSH2 domain form another important part of the autoinhibitory interface and two of these loops (β3/β4, β5/β6) within the C2 domain also showed heightened hydrogen-deuterium exchange. More peripheral regions of exchange include portions of the first two EF hands that directly support the β3/β4 loop of the C2 domain.

As described earlier, FGFR1K principally interacts with the nSH2 domain of PLC-γ1 (Bae et al., 2009; Chattopadhyay et al., 1999; Ji et al., 1999) such that many of the sites of increased exchange within PLC-γ1 are distant from the main binding site of FGFR1K. Consequently, while it is undoubtedly the case that the binding of FGFR1K to PLC-γ1 leads to widespread increases in the hydrogen-deuterium exchange kinetics of PLC-γ1, it is not readily apparent how FGFR1K promotes this increased exchange. Nonetheless, the increased exchange throughout the autoinhibitory interface is indicative of its increased mobility, suggesting that kinase engagement is sufficient to loosen the autoinhibitory interface and prime PLC-γ1 for more complete activation upon its phosphorylation by the kinase. This idea has been suggested previously (Hajicek et al., 2019).

In contrast to the widespread changes in the exchange profile of PLC-γ1 (H335A) upon engagement of phosphorylated FGFR1K, PIP_2_-containing liposomes incubated with PLC-γ1 (H335A) produced more limited differences that were mostly restricted to the periphery of the lipase (**Fig. 2B, S5**). Overall, there were no regions of significantly decreased exchange and only three, discrete, non-contiguous regions of significantly increased exchange. Two of the regions of increased exchange were restricted to relatively small portions of either the SH3 domain (β2) or the TIM barrel (β6/α5/α5’). A larger region of exchange manifested within the two SH2 domains and encompasses regions of the nSH2 domain (αA/βB) near the site of interaction with phosphorylated Tyr766 of FGFR1K, the linker between the two SH2 domains, and the terminal portion of the cSH2 domain (αB/βG) required for autoinhibition through interactions with the C2 domain. This third, larger region is compelling since it suggests a propensity of the nSH2 domain to interact with membranes and place it within proximity of phosphorylated receptors such as FGFR1 in order to facilitate the docking of PLC-γ1 with active transmembrane receptors. It also suggests that membranes impact regions between the two SH2 domains as well as within the cSH2 domain to possibly facilitate release of autoinhibition. Nonetheless, the overall magnitudes of differences in exchange in response to liposomes are small compared to the differences observed upon FGFR1K engagement (**Fig. 2A-B, S5**). This result suggests multiple points of transient engagement of PLC-γ1 with liposomes or cellular membranes in the absence of transmembrane binding partners. This scenario is also consistent with the tight, basal autoinhibition of PLC-γ1 in cells in the absence of a relevant, activated transmembrane receptor (Gresset et al., 2010). However, in the presence of relevant activated receptors as represented by phosphorylated FGFR1, the interaction of membranes with PLC-γ1 may help PLC-γ1 to engage receptors and may subsequently cooperate with receptors to facilitate release of lipase autoinhibition.

Cooperation between FGFR1 and membranes in the regulation of PLC-γ1 is also consistent with the hydrogen-deuterium exchange pattern of PLC-γ1 observed in the presence of both FGFR1K and liposomes (**Fig. 2C, S5**). Overall, similar patterns of differences in exchange are observed for the same regions of PLC-γ1 affected by either component alone. Indeed, the overlap of the exchange profiles is rather striking and is a testament to the reproducibility of the experimental technique. However, when both FGFR1K and liposomes were present, there was enhanced increases in exchange in PLC-γ1 compared to either condition alone. These increases manifest primarily within the SH3 domain and the catalytic TIM barrel. Overall, HDX-MS data suggest a modest, synergistic cooperation between FGFR1K and liposomes to increase the conformational flexibility of the regulatory domains relative to the catalytic core of PLC-γ1.

### Oncogenic substitution of PLC-γ1 mimics engagement by kinase and liposomes

The PLC-γ1 interface between its regulatory domains and catalytic core is frequently mutated in certain cancers and immune disorders (Koss et al., 2014). We have previously proposed that these mutations likely disrupt this interface leading to elevated, basal phospholipase activity (Hajicek et al., 2019). If true, oncogenic substitutions at this interface in PLC-γ1 might recapitulate changes in deuterium exchange in PLC-γ1 observed upon binding phosphorylated FGFR1K. This idea was tested using HDX-MS and PLC-γ1 (H335A) harboring an additional substitution (D1165H) frequently found in leukemias and lymphomas (Kataoka et al., 2015) (**Fig. 3**). Asp1165 resides in a loop between strands β5 and β6 of the C2 domain where it is required to stabilize extensive interactions between the C2 and cSH2 domains. These interactions are required to maintain PLC-γ1 in an autoinhibited state since the basal activity of PLC-γ1 (D1165H) is ~1,500-fold higher than the wild-type phospholipase (Hajicek et al., 2019).

**Figure 3.**
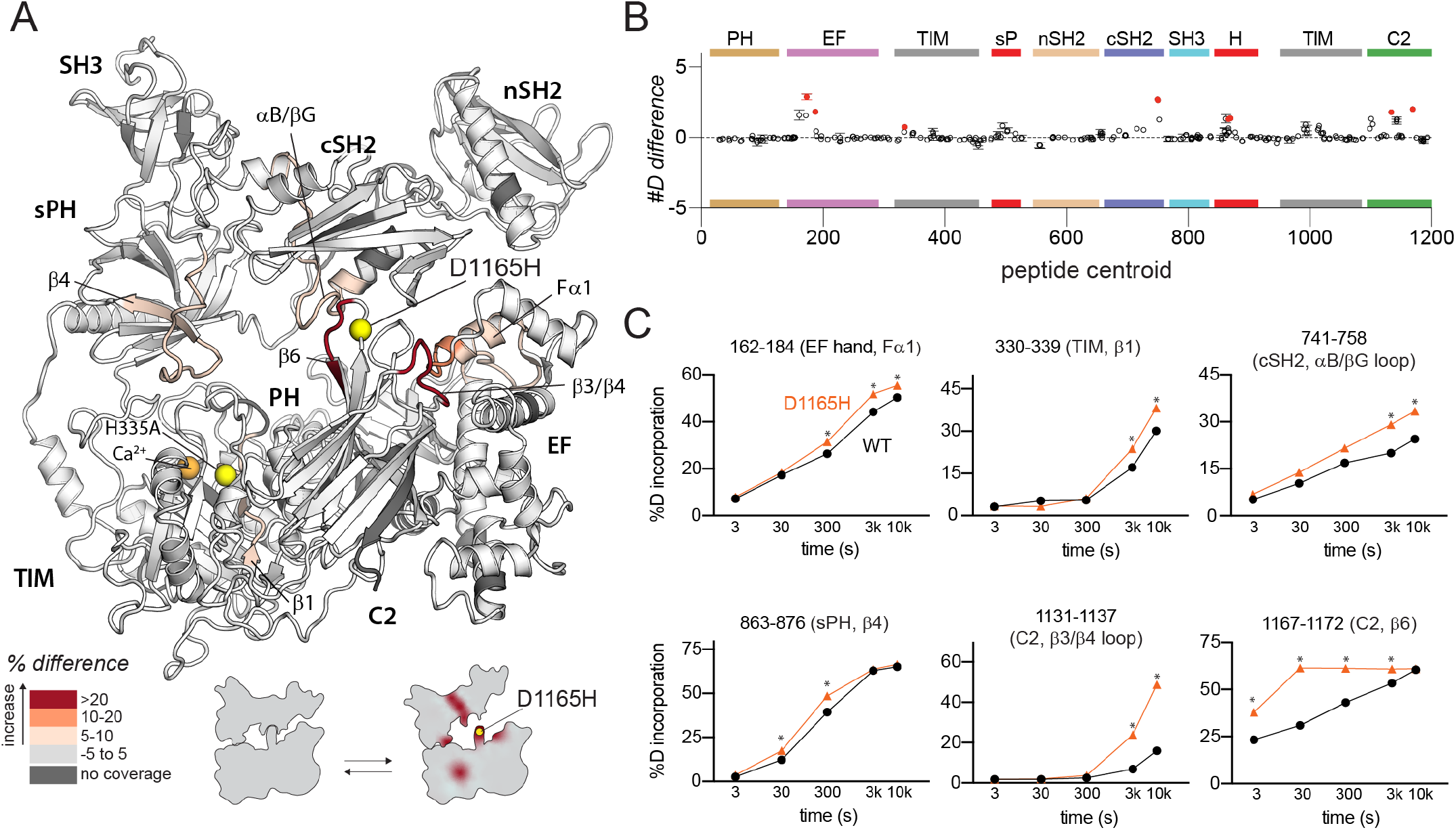
Oncogenic substitution of PLC-γ1 mimics kinase engagement. (**A**) Significant differences in deuterium incorporation for wild type PLC-γ1 (H335A) versus the oncogenic mutant of PLC-γ1 (H335A+D1165H) was determined and mapped on the structure of PLC-γ1. (**B**) The #D difference upon mutation of D1165H show the total difference in deuterium incorporation over the entire HDX time course, with each point indicating a single peptide (error shown as SD [n=3]). Individual peptides with a significant difference between conditions (defined as greater than both a 5% and 0.4 Da difference in exchange at any time point, and a two-tailed unpaired t-test of p<0.01) are colored red. (**C**) A selection of peptides showing significant differential exchange at any time point (data shown as mean ± SD [n = 3]), asterisks indicate significant times points for peptides as defined above. Full deuterium exchange data is available in the source data.

In comparison to catalytically-inactive PLC-γ1 harboring H335A, the same version of PLC-γ1 with the addition of D1165H exhibited increased deuterium exchange throughout the interface between the regulatory domains and the catalytic core (**Fig. 3**). In the immediate vicinity of D1165H, deuterium exchange increased within the β6 strand of the C2 domain suggesting that Asp1165 serves to support the position of the β6 strand. This idea is consistent with previous molecular dynamics simulations of PLC-γ1 that highlighted an unravelling of the β6 strand upon introduction of D1165H (Hajicek et al., 2019). The previous molecular dynamics simulations also highlighted a relatively large (~10 Å) movement of the cSH2 upon collapse of the β6 strand, and this movement may be reflected in the increased exchange of the nearby αB helix and αB/βG loop of the cSH2 domain. Other regions of increased exchange are more distant and include portions of the first EF hand, sPH domain, and TIM barrel. Overall, regions of increased exchange in catalytically-inactive PLC-γ1 (D1165H) overlap well with regions of increased exchange in PLC-γ1 (H335A) when it engages phosphorylated FGFR1K (**Fig. 2A**). This comparison further supports the notion that disengagement of the catalytic core from the regulatory domains will enhance lipase activity, with oncogenic mutations and binding to phosphorylated FGFR1K mediating this process. Indeed, it should be noted that regions of increased exchange are more extensive in the complex of PLC-γ1 and FGFR1K suggesting that oncogenic mutations cannot fully recapitulate activation by phosphorylated FGFR1K.

### Oncogenic substitution uncovers functional cooperativity within PLC-γ1

Constitutively-active variants of PLC-γ1 and -γ2 have untapped reserves of lipase activity. For example, although PLC-γ1 (D1165H) possesses exceptionally high basal lipase activity, this activity can be further enhanced in cells by the epidermal growth factor receptor suggesting that normal cellular regulation is partially preserved in constitutively active forms of PLC-γ1 (Hajicek et al., 2019). To test this idea, changes in the hydrogen-deuterium exchange profile of PLC-γ1 (H335A+D1165H) were determined in response to phosphorylated FGFR1K, liposomes, or both (**Fig 4, S6, S7**). In comparison to the identical measurements described earlier using PLC-γ1 (H335A), these measurements are particularly revealing for several reasons.

**Figure 4.**
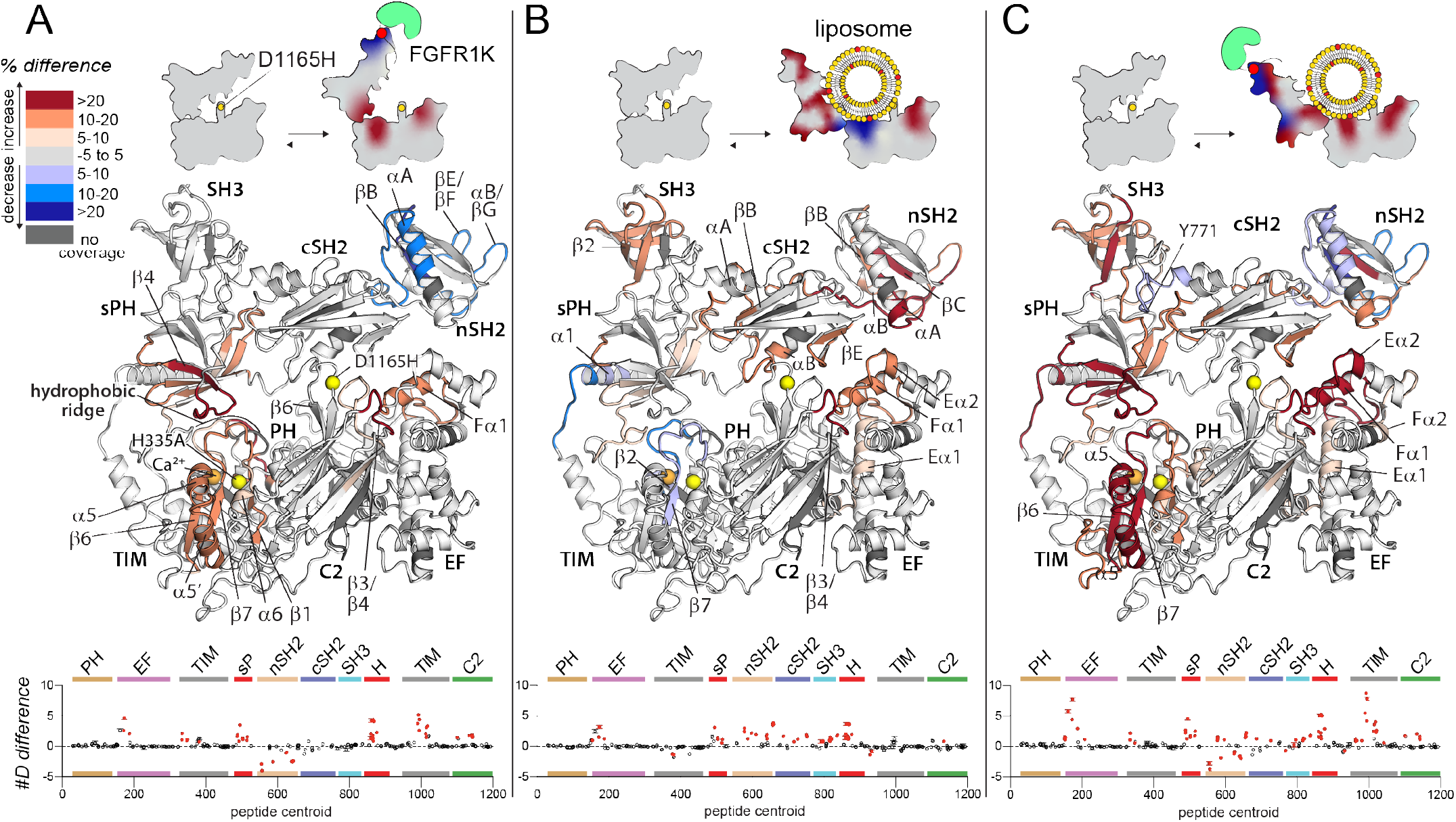
Oncogenic substitution uncovers functional cooperativity within PLC-γ1. Differences in deuterium incorporation were measured for PLC-γ1 (H335A+D1165H) in three states: bound to phosphorylated FGFR1K (**A**), in the presence of PE:PIP_2_ (90:10) liposomes (**B**), or with both kinase and liposomes (**C**). Differences in deuterium exchange for a series of time points (3, 30, 300, 3000, and 10000 sec) were calculated relative to PLC-γ1 (H335A+D1165H) alone and peptides with significant differential exchange at any time point (both a 5% and 0.4 Da difference in exchange at any time point, and a two-tailed unpaired t-test of p<0.01) mapped onto the structure. The #D difference for each condition is graphed below, which show the total difference in deuterium incorporation over the entire HDX time course, with each point indicating a single peptide (error shown as SD [n=3]); peptides with significant differences between conditions as defined above are red.

First, phosphorylated FGFR1K produced almost identical changes in the exchange profiles of the two forms of PLC-γ1 as highlighted in the comparison of figures 2A and 4A (see also **Fig. S6A**). Not only does this result further attest to the reproducibility of the experimental design, it also strongly suggests that wild-type and constitutively active forms of PLC-γ1 respond similarly to engagement by kinases.

Second, and in contrast to effects mediated by phosphorylated FGFR1K, PIP_2_-containing liposomes produced much more extensive and wide-spread changes in the hydrogen-deuterium exchange profile of mutant PLC-γ1 (H335A+D1165H) (**Figs. 4B, S6B**) compared to PLC-γ1 (H335A) (**Fig. 2B**). For example, while no portion of the EF hands of PLC-γ1 (H335A) showed differences in exchange upon addition of liposomes, the entire first EF hand (Eα1/Fα1) as well as the start of the second EF hand (Eα2) of PLC-γ1 (H335A+D1165H) showed increased exchange. In addition, the adjacent β3/β4 loop of the C2 domain had robustly increased exchange upon membrane binding in the D1165H variant, whereas this region in PLC-γ1 (H335A) was essentially unresponsive to lipids. In the same vein, while liposomes elicited only relatively modest and discrete increases in the exchange of the regulatory domains of PLC-γ1 (H335A), under the same conditions, there were widespread and robust increases in exchange within the regulatory domains of PLC-γ1 (H335A+D1165A). These regions include the entire first half of the sPH domain, the majority of both the nSH2 and SH3 domains, as well as the EF and BG loops (βE/βF, αB/βG) of the cSH2 domain critical for autoinhibition of lipase activity.

Third, and perhaps most intriguing, while liposomes failed to decrease exchange of any part of PLC-γ1 (H335A), under identical conditions, several portions of PLC-γ1 (H335A+D1165H) exhibited decreased exchange. These regions of decreased exchange are primarily in the TIM barrel: β2-β2’ and β7 strands as well as the subsequent loops comprising the majority of the hydrophobic ridge. The hydrophobic ridge is generally assumed to insert into membranes during interfacial catalysis and the decreased exchange of this region may indicate its direct interaction with liposomes during aborted attempts at catalysis. A second region of decreased exchange encompasses α1 of the sPH domain as well as the adjoining linker connecting to the second half of the TIM barrel. This linker is disorder in the crystal structure of full-length PLC-γ1 and it is intriguing to speculate that this region may act as a hinge that allows the regulatory array to move in order to allow access of liposomes to the hydrophobic ridge.

Finally, the exchange profile of PLC-γ1 (H335A+D1165H) upon the addition of both phosphorylated FGFR1K and liposomes (**Fig. 4C**) approximately resembles the composite profile of each component alone with two interesting caveats. First, a portion of the linker between the cSH2 and SH3 domains that includes frequently phosphorylated Tyr771 possesses decreased exchange that cannot be accounted by a composite profile. This difference presumably reflects structural changes in PLC-γ1 (H335A+D1165H) that arise due to cooperative interactions among the PLC isozyme, FGFR1K, and liposomes. This presumption is reinforced by the second caveat: in the presences of both FGFR1K and liposomes, there is a striking synergy in the differences in exchange for PLC-γ1 (H335A+D1165H) compared to PLC-γ1 (H335A), with liposomes enhancing the changes observed with only FGFR1K approximately 2-fold (**Fig. 4C, lower panel**). These regions include most of the first and second EF hands as well as a large segment of the second half of the TIM barrel spanning strands β6 and β7 and including helices α5 and α5’.

Overall, the exchange results indicate that the oncogenic substitution, D1165H, leads to a much more conformationally-flexible PLC-γ1 that more readily interacts with lipid bilayers, possibly in a fashion that mirrors interfacial catalysis by PLC isozymes. This process would require a large movement by the regulatory domains as suggested by the icons in Figure 4.

### Phosphorylated FGFR1K increases PLC-γ1 specific activity

The hydrogen-deuterium exchange kinetics described above strongly suggest that the initial engagement of PLC-γ isozymes by kinases may be sufficient to enhance phospholipase activity irrespective of subsequent phosphorylation. This idea was tested using XY-69 (Huang et al., 2018), a fluorescent analog of PIP_2_, embedded in lipid vesicles. (**Fig. 5**). PLCs readily hydrolyze XY-69 leading to increased fluorescence that can be followed in real-time. As shown previously, when XY-69 in lipid vesicles is mixed with PLC-γ1, the specific activity of the phospholipase is low, indicative of basal autoinhibition. This activity remained unchanged upon pre-incubation of PLC-γ1 with catalytically-inactive FGFR1K that was not phosphorylated. In contrast, when PLC-γ1 was pre-incubated with catalytically-inactive FGFR1K phosphorylated at Tyr766 to promote complex formation with PLC-γ1, the specific lipase activity of PLC-γ1 increased approximately three-fold. These results indicate that kinase engagement by PLC-γ1 is sufficient to enhance lipase activity and supports the proposition that PLC-γ1 is allosterically modulated by FGFR1K upon complex formation and irrespective of the phosphorylation state of the lipase.

**Figure 5.**
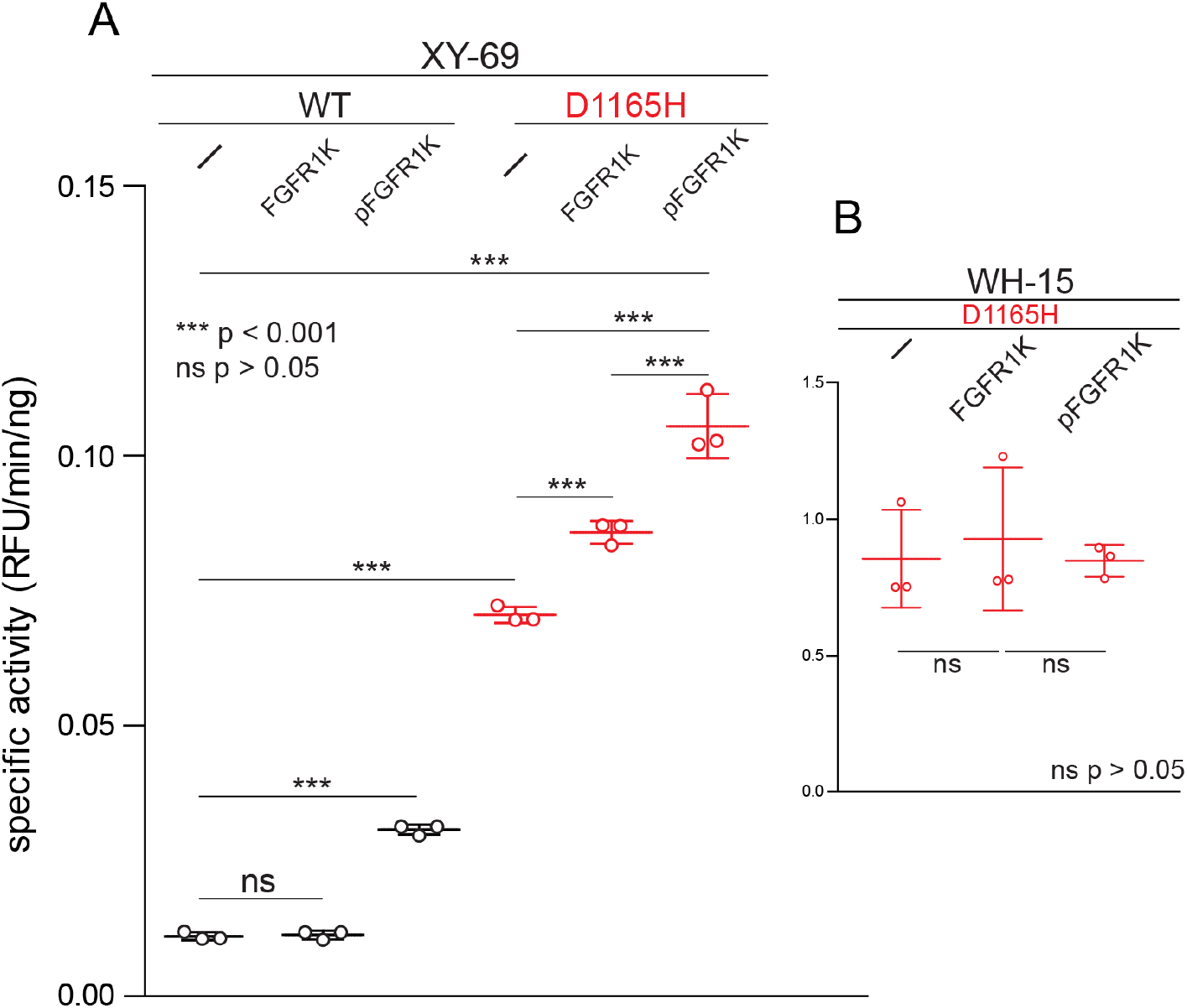
Phosphorylated FGFR1K increases PLC-γ1 specific activity. (**A**) Specific activities measured with the membrane-embedded substrate XY-69. XY-69 (0.5μM) was incorporated into liposomes comprised of PE:PIP_2_ (80:20) prior to addition of indicated variants of PLC-γ1 (10 nM WT; 0.3 nM D1165H). Specific activities were calculated for PLC-γ1 alone and in the presence of either unphosphorylated FGFR1K or its phosphorylated counterpart (pFGFR1K). (**B**) Specific activities of PLC-γ1 (D1165H) measured with water-soluble WH-15 (5 μM). In all cases, specific activities are presented as the mean ± SD of three independent experiments (n=3), each with three or more technical replicates. Statistical significance was determined with a one-way ANOVA followed by Tukey’s post-hoc test.

Similar results were observed with PLC-γ1 (D1165H). In particular, addition of catalytically-inactive, phosphorylated FGFR1K increased the basal lipase activity of PLC-γ1 (D1165H) approximately two-fold. The basal capacity of PLC-γ1 (D1165H) to hydrolyze vesicle-embedded XY-69 was substantially higher than the equivalent activity of wild-type PLC-γ1, but this result has been previously published (Huang et al., 2018) and comports with the higher constitutive activity of PLC-γ1 (D1165H) in cells (Patel et al., 2020). Interestingly, the addition of catalytically-inactive but non-phosphorylated FGFR1K also led to a significant increase in the specific activity of PLC-γ1 (D1165H). We are unable to readily account for this observation, but it should be noted that the crystal structure of phosphorylated FGFR1K bound to the two SH2 domains of PLC-γ1 highlights a secondary interface outside of the canonical pTyr-binding site that contributes to the affinity of the complex (Bae et al., 2009). Therefore, unphosphorylated FGFR1K may have sufficient affinity for PLC-γ1 that it is able to enhance the lipase activity of constitutively active PLC-γ1 (D1165H) but not the more basally-autoinhibited, wild-type form.

Autoinhibition of the PLC-γ isozymes presumably arises due to steric hindrance by the regulatory array preventing the catalytic core from productively engaging lipid membranes to hydrolyze XY-69. In contrast, WH-15 is a soluble, fluorescent substrate of PLCs that is insensitive to the autoregulation of the PLC-γ isozymes (Huang et al., 2011). As expected, the hydrolysis of WH-15 by PLC-γ1 (D1165H) is insensitive to either form of FGFR1K (**Fig. 5B**) and these results further support the idea that the complex of FGFR1K and PLC-γ1 diminishes autoregulation by loosening the conformational ensemble of the autoinhibited form of PLC-γ1.

## Discussion

A general model for the regulated activation of the PLC-γ isozymes posits a conserved, catalytic core that is basally prevented from accessing its lipid substrate, PIP_2_, when PIP_2_ is embedded in membranes (**Fig. 6**). In essence, the twodimensional nature of the membrane bilayer is key to this regulation since soluble substrates such as PIP_2_ in detergent bypass this regulation and freely access the solvent-accessible active site. In contrast, the active site of PLC-γ isozymes is prevented from accessing PIP_2_ in membranes by a set of regulatory domains that must undergo conformational rearrangements before the catalytic core can dock with membranes and engage PIP_2_. This model derives from substantial cellular (DeBell et al., 2007; DeBell et al., 1999; Horstman et al., 1999; Poulin et al., 2000; Schade et al., 2016), biochemical (Bunney et al., 2012; Gresset et al., 2010; Poulin et al., 2005), and biophysical (Bunney et al., 2012; Hajicek et al., 2013) data, and is perhaps most self-evident for the high-resolution structure of autoinhibited, full-length PLC-γ1 that highlights the arrangement of the regulatory domains relative to the catalytic core that prevents productive engagement of membranes (Hajicek et al., 2019).

**Figure 6.**
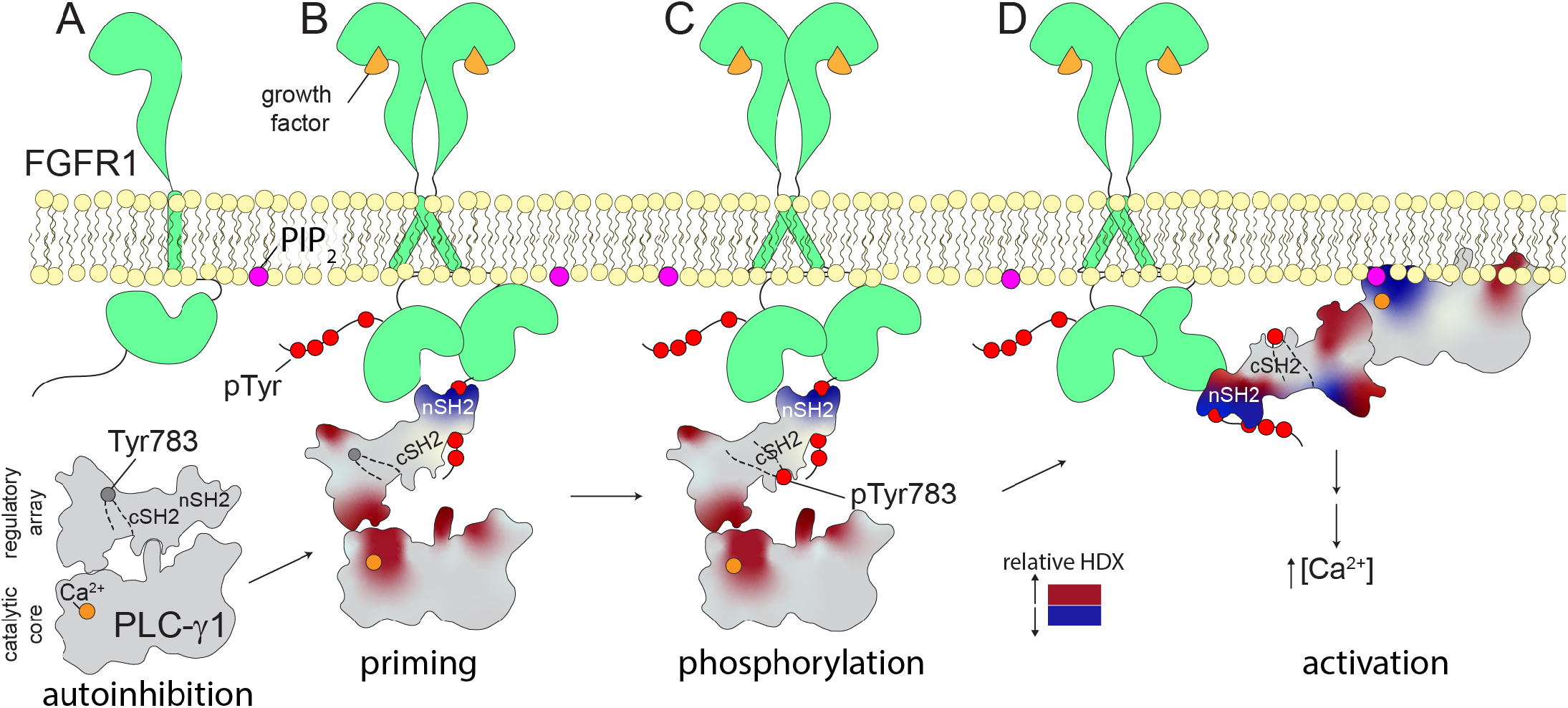
Multi-step activation of PLC-γ1. Initially, FGFR1 is monomeric and inactive while PLC-γ1 is cytosolic and basally autoinhibited (**A**). Growth factor-promoted dimerization of FGFR1 leads to its tyrosine phosphorylation and the recruitment of PLC-γ1 through interactions mediated by the nSH2 domain of PLC-γ1 (**B**). Complex formation reduces deuterium exchange within the nSH2 domain while simultaneously increasing deuterium exchange throughout the interface between the catalytic core and regulatory domains of PLC-γ1. Increased deuterium exchange is consistent with a loosening of the autoinhibited form of PLC-γ1 shown schematically. Active FGFR1 subsequently phosphorylates Tyr783 of PLC-γ1 to favor the interaction of pTyr783 and the cSH2 domain of PLC-γ1 (**C**). This step reinforces the disruption of the interface between regulatory and catalytic domains to favor a fully open form of PLC-γ1 capable of productively engaging membranes and manifesting as a decrease in the deuterium exchange of the catalytic TIM barrel (**D**).

What is much less well understood is the sequence of events culminating in the conformational rearrangements required for the activation of the PLC-γ isozymes. It is universally accepted that the phosphorylation of Tyr783 in PLC-γ1 or the equivalent residue, Tyr759, in PLC-γ2 is absolutely required for regulated activation, but the steps preceding or subsequent to these phosphorylations continue to be debated. The studies presented here address these less well understood steps and support the view of the PLC-γ isozymes as relatively quiescent lipases prior to recruitment to membranes. Indeed, a major conclusion of these studies is that recruitment to membranes appears to be integral to activation.

Two aspects of this recruitment and activation deserve special consideration. First is the engagement of PLC-γ isozymes with receptor tyrosine kinases or co-receptors. Complex formation requires receptors or co-receptors to be pre-phosphorylated at specific tyrosines to provide docking sites for the PLC-γ isozymes. Based on the majority of previous studies that address this point, the nSH2 domain of the PLC-γ isozymes directly engage the phosphorylate tyrosines in the myriad of relevant, membrane-resident receptors and co-receptors. This point is also consistent with the relatively large decreases in amide proton exchange of the nSH2 domain of PLC-γ1 upon addition of the phosphorylated kinase domain of FGFR1. However, much more interesting is the wide-spread increases in amide exchange of PLC-γ1 upon complex formation. These wide-spread increases highlight a phospholipase that is exquisitely—and indeed, globally—tuned to engagement of the kinase. The most significant implication being that engagement of the nSH2 domain by a phosphorylated receptor or co-receptor increases the dynamic flexibility of large portions of the phospholipase. This increased flexibility will tend to loosen the autoinhibited state of the phospholipase to promote activation. In essence, initial recruitment of the PLC-γ isozymes to phosphorylated receptors or co-receptors is a crucial first step for lipase activation. We previously inferred this “priming” step based solely on structural considerations, and the exchange data presented here strongly support this idea. Indeed, since priming is predicted to disfavor the autoinhibited state, it is also predicted to favor activation. Thus a priming step—that is independent of subsequent phosphorylation of PLC-γ1—readily explains the increased lipase activity of PLC-γ1 operating on membrane-embedded PIP_2_ and bound to the phosphorylated, but catalytically-inactive, kinase domain of FGFR1.

The second aspect to consider is the intrinsic role of membranes during the activation of PLC-γ isozymes. Lipid vesicles containing PIP_2_ were relatively ineffectual in altering the amide exchange of wild-type PLC-γ1. This situation comports with the tight basal autoinhibition of PLC-γ isozymes in cells where the isozymes are inevitably in close proximity to lipid membranes. In contrast, the exchange kinetics of wild-type PLC-γ1 bound to the phosphorylated kinase domain of FGFR1 were amplified by the addition of lipid vesicles. Amplified exchange was even more extreme for PLC-γ1 (D1165H) containing a substitution introduced to perturb the interface between the catalytic core and regulatory domains (Hajicek et al., 2019). These results indicate that once primed by kinase—or indeed, mutation—PLC-γ1 becomes more sensitive to the presence of lipid vesicles. The net outcome likely shifts PLC-γ1 toward conformations that can productively engage membranes to hydrolyze membrane-bound PIP_2_.

These two aspects should not be considered in isolation. That is, the activation of PLC-γ isozymes involves the intertwined events of: i) docking to membrane-resident receptors or co-receptors and ii) the forced proximity at membranes of the dock complexes. The two events cooperate to initiate activation of PLC-γ isozymes (**Fig. 6**), presumably by increasing accessibility to membrane-bound PIP_2_. At this point, it should be emphasized that the experimental setup studied here does not fully recapitulate the cellular activation process and almost certainly underestimates the degree of cooperativity. More specifically, to facilitate experimental tractability, and especially the desire to study complex formation independent of recruitment to membranes, these studies exclusively used an isolated kinase domain of FGFR1 *not* embedded in membranes. We predict cooperativity will be greatly enhanced for receptors or co-receptors embedded in membranes as typically found in cells. Increased cooperativity will result in increased sensitivity of activation of the PLC-γ isozymes. It should also be noted that these two steps along the activation pathway are general in that they are likely to manifest for any receptor or co-receptor that recruits PLC-γ1 or -γ2 to membranes through motifs that contain phosphorylated tyrosine and bind the nSH2 domain of PLC-γ isozymes.

Up to this point, we have considered only membrane recruitment and activation of PLC-γ isozymes by phosphorylated receptors and coreceptors. However, these are not the only proteins that bind PLC-γ isozymes at membranes. For example, several scaffolding proteins such as SLP-76 that contain polyproline regions bind the SH3 domain of PLC-γ isozymes (Kim et al., 2000; Manna et al., 2018; Rouquette-Jazdanian et al., 2012; Tvorogov and Carpenter, 2002). Similarly, the small GTPase, Rac2, specifically engages the sPH domain of PLC-γ2 to promote lipase activation (Bunney et al., 2009; Piechulek et al., 2005; Walliser et al., 2008). We do not know if these complexes affect the conformational dynamics of the PLC-γ isozymes. However, if they do, these complexes may also prime the PLC-γ isozymes for activation at membranes with one potential outcome being an intricate interplay of multiple binding events regulating the activation of the PLC-γ isozymes.

## Material and Methods

### Molecular cloning

#### PLC-γ1

Transfer vector (pFB-LIC2) encoding wild-type rat PLC-γ1 (residues 21-1215) and PLC-γ1 (D1165H) have been previously described. The H335A mutation that renders PLC-γ1 catalytically inactive was introduced into the transfer vector encoding wild-type PLC-γ1 (residues 21-1215) or PLC-γ1 (D1165H) using standard primer-mediated mutagenesis (Agilent Technologies; QuikChange site-directed mutagenesis manual). Mutagenesis was confirmed by automated DNA sequencing of the open reading frame. The PLCs encoded by these constructs are referred as PLC-γ1 (H335A) and PLC-γ1 (H335A+D1165H), respectively.

#### FGFR1K

Plasmid DNA encoding the kinase domain (residues 458-774) of human fibroblast growth factor receptor 1 (FGFR1) with four substitutions (Y463F, Y583F, Y585F, Y730F) that eliminate sites of phosphorylation was synthesized by Genewiz (South Plainfield, NJ). The plasmid DNA encoding this variant of FGFR1K was then sub cloned from a pUC57-Amp vector to a modified pFastBac-HT vector (pFB-LIC2) using a ligation-independent cloning (LIC) strategy. The baculovirus expression vector, pFB-LIC2, incorporates a His6 tag and a tobacco etch virus (TEV) protease site at the amino terminus of the expressed protein. This transfer vector was then mutated to encode two forms of FGFR1K: i) a version mutated (Y653F and Y654F) to prevent phosphorylation-dependent activation of kinase activity, i.e., kinase inactive, and ii) a version that retained kinase activity but was mutated (Y766F) to prevent phosphorylation-dependent interaction with PLC-γ1. Mutations were introduced using standard primer-mediated mutagenesis (Agilent Technologies; QuikChange site-directed mutagenesis manual) and all FGFR1K variants were verified by automated DNA sequencing of the corresponding open reading frames.

### Protein expression and purification

#### PLC-γ1

Expression and purification of PLC-γ1 variants was previously described and used here with minor modifications. Briefly, four liters of High Five (*T. ni*) cells at a density of approximately 2.0 x 10^6^ cells/mL were infected with amplified baculovirus stock (10 - 15 mL/L) encoding individual PLC-γ1 variants. Cells were harvested approximately 48 hr. post-infection by centrifugation at 6,000 rpm in a Beckman JA-10 rotor at 4°C. Cell pellet was resuspended in 200 mL of buffer N1 (20 mM HEPES (pH 7.5), 300 mM NaCl, 10 mM imidazole, 10% v/v glycerol, and 0.1 mM EDTA) supplemented with 10 mM 2-mercaptoethanol (BME) and four EDTA-free cOmplete protease inhibitor tablets (Roche Applied Science) prior to lysis using the Nano DeBEE High Pressure Homogenizer (BEE International). Lysate was centrifuged at 50,000 rpm for 1 hr. in a Beckman Ti70 rotor. The supernatant was filtered through a 0.45 μm polyethersulfone (PES) low protein-binding filter and loaded onto a 5 mL HisTrap HP immobilized metal affinity chromatography (IMAC) column equilibrated in buffer N1. The column was washed with 15 column volumes (CV) of buffer N1, followed by 15 CV of 2.5% buffer N2 (buffer N1 + 1 M imidazole). Bound proteins were eluted with 40% buffer N2. Fractions containing PLC-γ1 were pooled and dialyzed overnight in the presence of 2% (w/w) TEV protease to remove the His6 tag in a buffer solution containing 20 mM HEPES (pH 7.5), 300 mM NaCl, 10% v/v glycerol, 1 mM dithiothreitol (DTT), 1 mM EDTA. The sample was subsequently diluted 2-fold with buffer N1 and applied to a 5 mL HisTrap HP column. Flow-through fractions containing cleaved PLC-γ1 were pooled, diluted 4-fold with buffer Q1 (20 mM HEPES (pH 7.5) and 2 mM DTT), and loaded onto an 8 mL SourceQ anion exchange column equilibrated in 10% buffer Q2 (buffer Q1 + 1 M NaCl). Bound proteins were eluted in a linear gradient of 10% - 60% buffer Q2 over 50 CV. Fractions containing PLC-γ1 were pooled, concentrated using a GE Healthcare VivaSpin 50K molecular weight cut-off (MWCO) centrifugal concentrator and applied to a 16 mm x 700 mm HiLoad Superdex 200 size exclusion column equilibrated in a buffer solution containing 20 mM HEPES (pH 7.5), 150 mM NaCl, 5% v/v glycerol, and 2 mM DTT. Pure PLC-γ1 was concentrated to a final concentration of 40 - 80 mg/mL, aliquoted, snap-frozen in liquid nitrogen, and stored at −80°C until use.

#### FGFR1K

Expression and purification of kinase-inactive FGFR1K follows that of the PLC-γ1 variants with the following modifications. After removal of the His6 tag by TEV protease and subsequent 5 mL HisTrap HP IMAC column, the sample was concentrated using a GE Healthcare VivaSpin 10K MWCO centrifugal concentrator. Concentrated sample was applied to a 16 mm x 700 mm HiLoad Superdex 200 size exclusion column equilibrated in buffer containing 20 mM HEPES (pH 7.5), 200 mM NaCl, 5% (v/v) glycerol, and 2 mM DTT. Fractions containing pure, kinase-inactive FGFR1K were concentrated to a final concentration of 20 - 30 mg/mL, aliquoted, snap-frozen in liquid nitrogen, and stored at −80°C until use. The kinase active from of FGFR1K harboring Y766F was expressed as described above, however, purification terminated after the first 5 mL HisTrap HP IMAC column so the protein retains its His6 tag.

### In vitro phosphorylation of FGFR1K

Equimolar concentrations (100 μM) of tagless, kinase-inactive FGFR1K and the tagged, kinase-active version were incubated in 20 mM HEPES (pH 7.5), 50 mM NaCl, 25 mM MgCl_2_, 50 ng/mL fatty-acid free bovine serum albumin (FAF BSA), 10 mM ATP, and 2 mM DTT. The phosphorylation reaction was terminated after 100 minutes by adding EDTA to a final concentration of 50 mM and kinase phosphorylation was confirmed via native PAGE followed by staining with Coomassie Brilliant Blue. To separate the two forms of FGFR1K, the mixture was loaded onto a 1 mL HisTrap HP IMAC using a 200 μL sample loop followed by 2 mL of buffer N1 (20 mM HEPES pH 7.5, 100 mM NaCl, 10 mM MgCl_2_, and 2 mM DTT). The column was washed with 5 CV of buffer N1, followed by 5 CV of 40% buffer N2 (buffer N1 + 1 M imidazole), and 5 CV of 100% buffer N2. Fractions containing phosphorylated, kinase-inactive FGFR1K were pooled and concentrated using a GE Healthcare Vivaspin 6 10 kDa MWCO, aliquoted, snap-frozen in liquid nitrogen, and stored at −80°C until use.

### LC-MS/MS of phosphorylated FGFR1K

Phosphorylated, kinase-inactive FGFR1K was diluted and loaded onto native PAGE followed by staining with Coomassie Brilliant Blue. Gel bands were excised and digested with trypsin overnight. Peptides were extracted, then analyzed by LC/MS/MS using the Thermo Easy nLC 1200-QExactive HF. Data were searched against a UniProt Sf9 database including the sequence for FGFR1K using Sequest within Proteome Discoverer 2.1. All data were filtered using a false discovery rate of 5%.

### Formation of PLC-γ1 in complex with FGFR1K

Kinase-inactive FGFR1K phosphorylated at Tyr766 was generated as described above. Either PLC-γ1 (H335A) or PLC-γ1(H335A+D1165H) was added in a 2-fold molar excess relative to kinase. The sample was immediately loaded onto a 10 mm × 300 mm Superdex 200 GL size exclusion column equilibrated with 20 mM HEPES (pH 7.5), 100 mM NaCl, 10 mM MgCl_2_, and 2 mM DTT. Pure complex was pooled and concentrated using a GE Healthcare Vivaspin 6 10K MWCO, aliquoted, snap-frozen in liquid nitrogen, and stored at −80°C until use. Formation of the complex was confirmed via SDS-PAGE followed by staining with Coomassie Brilliant Blue.

### In vitro quantification of phospholipase activity

#### XY-69 fluorogenic assay

Liposomes with a final PE:PIP_2_ content of 80:20 were generated by mixing 750 nM XY-69 (24), 192 μM of liver phosphatidylethanolamine (PE, Avanti Polar Lipids), 48 μM brain phosphatidylinositol 4,5-bisphosphate (PIP_2_, Avanti Polar Lipids) in 12 x 75 mm borosilicate tubes. Lipids were dried under a nitrogen stream followed by high vacuum (0.5 mtorr). Dried lipid mixture was suspended in 20 mM HEPES (pH 7.5) using a probe microtip sonicator of 5/64” at 20% output for 3 cycles of 5 sec ON, 15 sec OFF. Concurrently, the PLC-γ1 proteins were diluted in a buffer containing 20 mM HEPES (pH 7.5), 50 mM NaCl, 2 mM DTT, and 1 mg/mL FAF BSA. The 6X assay buffer containing 80 mM HEPES (pH 7.5), 420 mM KCl, 10 mM DTT, 18 mM EGTA, and 14.1 mM CaCl_2_ (~390 nM free Ca^2+^) was added to the resuspended lipid mixture in a 1:4 ratio. To a non-binding surface (NBS)-treated Corning 384-well plate, 2 μL of diluted PLC-γ1 proteins were added, either alone or in the presence of a two-fold molar excess of unphosphorylated or phosphorylated, kinase-inactive FGFR1K relative to PLC-γ1. To initiate the assay, 10 μL of the lipid and assay buffer mixture was added. The plates were incubated at 30°C and data was recorded for 30 min at intervals of 1 min using excitation and emission wavelengths of 485 nm and 520 nm, respectively. Fluorescence intensity was normalized to a blank reaction lacking phospholipase, and initial rates of XY-69 hydrolysis were calculated from the slope of the linear portion of the curve. Final concentrations of XY-69, PIP_2_, and PE were 5 μM, 48 μM, and 192 μM, respectively.

#### WH-15 fluorogenic assay

Assays with the soluble substrate WH-15 were performed as described previously (25) with the following modifications. WH-15 was diluted to a final concentration of 5 μM in assay buffer containing 50 mM HEPES (pH 7.5), 70 mM KCl, 3 mM EGTA, 2.97 mM CaCl_2_ (~10 μM free Ca^2+^), 50 μg/mL FAF BSA, and 2 mM DTT. Basal fluorescence was equilibrated for approximately 10 minutes before addition of PLC-γ1 (D1165) alone or in the presence of a two-fold molar excess of unphosphorylated or phosphorylated, kinase-inactive FGFR1K relative to PLC-γ1 (1nM, final concentration). Data were recorded for 1 hr at intervals of 30 sec using excitation and emission wavelengths of 488 nm and 520 nm, respectively. Fluorescence intensity was normalized to the reaction lacking phospholipase, and initial rates of WH-15 hydrolysis were calculated from the slope of the linear portion of the curve.

### Liposome floatation assay

Liposomes with a final content of 89.8% PE, 10% PIP_2_, and 0.2% 7-nitro-2-1,3-benzoxadiazol-4-yl (NBD)-PE were prepared as described above and resuspended in 20 mM HEPES (pH 7.5). Concurrently PLC-γ1 (H335A), alone or in complex with phosphorylated, kinase-inactive FGFR1K were diluted in assay buffer containing 100 mM HEPES (pH 7.5), 150 mM NaCl, 10 mM KCl, 10 μM CaCl_2_, 0.5 mM tris(2-carboxyethyl)phosphine (TCEP). PLC-γ1 and liposomes were mixed and incubated on ice for 2 min. Immediately after mixing, 100 μL of assay buffer with 75% sucrose were added to the protein-lipid mixture to a final concentration of 30% sucrose in 250 μL total volume. The samples were overlaid with 200 μL of assay buffer with 25% sucrose and 50 μL of assay buffer. Samples were centrifuged at 55,000 rpm in a TLS-55 rotor for 1 hr at 4°C. After centrifugation, three fractions were collected: bottom (B, 250 μL), middle (M, 150 μL), and top (T, 100 μL). Samples (30 μL) of each fraction were mixed with 7 μL of 6X SDS sample buffer and analyzed by SDS-PAGE followed by staining with Coomassie Brilliant Blue.

### Hydrogen deuterium exchange-mass spectrometry

#### Sample preparation

Exchange reactions were carried out at 18°C in 20 μL volumes with a final concentration of 1.25 μM PLC-γ1 (H335A), PLC-γ1 (H335A+D1165H), PLC-γ1 (H335A)-FGFR1K complex, or PLC-γ1 (H335A+D1165H)-FGFR1K complex. A total of eight conditions were assessed: four in the presence of liposomes containing 90% phosphatidylethanolamine (PE) and 10% phosphatidylinositol 4,5-bisphosphate (PIP_2_) and four in the absence of liposomes. The conditions were as follows:

PLC-γ 1(H335A) alone
PLC-γ 1(H335A+D1165H) alone
PLC-γ1(H335A)-FGFR1K
PLC-γ1(H335A+D1165H)-FGFR1K
PLC-γ1(H335A) + liposomes
PLC-γ 1(H335A+D1165H) + liposomes
PLC-γ1(H335A)-FGFR1K + liposomes
PLC-γ1(H335A+D1165H)-FGFR1K + liposomes

For conditions containing liposomes, lipids were present at a final concentration of 320 μM. Prior to the addition of D_2_O, 2.0 μL of liposomes (or liposome buffer (20 mM HEPES pH 7.5, 100 mM KCl)) were added to 1.5 μL of protein, and the solution was left to incubate at 18°C for 2 min. Hydrogen-deuterium exchanges was initiated by the addition of 16.5 μL D_2_O buffer (88% D_2_O, 150 mM NaCl, 100 mM HEPES pH 7.0, 10 μM CaCl_2_, 0.5 mM TCEP pH 7.5) to 3.5 μL protein or protein-liposome solution for a final D2O concentration of 72%. Exchange was carried out over five time points (3, 30, 300, 3000, and 10000 sec) and terminated by the addition of 50 μL ice-cold acidic quench buffer (0.8 M guanidine-HCl, 1.2% formic acid). After quenching, samples were immediately frozen in liquid nitrogen and stored at −80°C. All reactions were carried out in triplicate.

#### Protein digestion and MS/MS data collection

Protein samples were rapidly thawed and injected onto an integrated fluidics system containing a HDx-3 PAL liquid handling robot and climate-controlled chromatography system (LEAP Technologies), a Dionex Ultimate 3000 UHPLC system, as well as an Impact HD QTOF Mass spectrometer (Bruker). Proteins were run over two immobilized pepsin columns (Applied Biosystems; Poroszyme™ Immobilized Pepsin Cartridge, 2.1 mm x 30 mm; Thermo-Fisher 2-3131-00; at 10°C and 2°C respectively) at 200 μL/min for 3 minutes. The resulting peptides were collected and desalted on a C18 trap column (Acquity UPLC^®^ BEH™ C18 1.7μm column (2.1 x 5 mm); Waters 186002350). The trap was subsequently eluted in line with a C18 reverse-phase separation column (Acquity 1.7 μm particle, 100 × 1 mm^2^ C18 UPLC column, Waters 186002352), using a gradient of 5-36% B (Buffer A 0.1% formic acid; Buffer B 100% acetonitrile) over 16 minutes. Full details of the LC setup and gradient are (Stariha et al., 2021). Mass spectrometry experiments were performed on an Impact II QTOF (Bruker) acquiring over a mass range from 150 to 2200 *m/z* using an electrospray ionization source operated at a temperature of 200 °C and a spray voltage of 4.5 kV.

#### Peptide identification

Peptides were identified using data-dependent acquisition following tandem MS/MS experiments (0.5 s precursor scan from 150-2000 m/z; twelve 0.25 s fragment scans from 150-2000 m/z). MS/MS datasets were analyzed using PEAKS7 (PEAKS), and a false discovery rate was set at 1% using a database of purified proteins and known contaminants (Dobbs et al., 2020).

#### Mass analysis of peptide centroids and measurement of deuterium incorporation

HD-Examiner Software (Sierra Analytics) was used to automatically calculate the level of deuterium incorporation into each peptide. All peptides were manually inspected for correct charge state, correct retention time, and appropriate selection of isotopic distribution. Deuteration levels were calculated using the centroid of the experimental isotope clusters. Results for these proteins are presented as relative levels of deuterium incorporation and the only control for back exchange was the level of deuterium present in the buffer (72%). Changes in any peptide at any time point (3, 30, 300, 3000, and 10,000 seconds) greater than specified cut-offs (5% and 0.4 Da) and with an unpaired, two-tailed t-test value of p<0.01 were considered significant.

The raw peptide deuterium incorporation graphs for a selection of peptides with significant differences are shown (**Fig. 3, S5, S6**), with the raw data for all analyzed peptides in the source Excel file. To allow for visualization of differences across all peptides, we used number of deuteron difference (#D) plots (**Fig. 2, 3, 4, S5, S6, S7**). These plots show the total difference in deuterium incorporation over the entire HDX time course, with each point indicating a single peptide. These graphs are calculated by summing the differences at every time point for each peptide and propagating the error (for example, **Fig. 2A**). For a selection of peptides, we are showing the %D incorporation over a time course, which allows for comparison of multiple conditions at the same time for a given region (**Fig. S5C, 6C**). Samples were only compared within a single experiment and were never compared to experiments completed at a different time with a different final D_2_O level. The data analysis statistics for all HDX-MS experiments are in **Table S1** and conform to recommended guidelines (Masson et al., 2019). The MS proteomics data have been deposited to the ProteomeXchange Consortium via the PRIDE partner repository (Perez-Riverol et al., 2019) with the dataset identifier PXD030492.

## Data availability

All HDX-MS mass spectrometry data in the paper are available from the PRIDE database with the accession number PXD030492. All raw H/D exchange data are available in the source data.

## Acknowledgements

We acknowledge members of the Sondek and Burke labs for their valuable feedback and technical support. We also thank Laura Herring and Josh Beri at the UNC Michael Hooker Proteomics Center for assistance with LC-MS/MS of phosphorylated kinase.

## Funding and additional information

This work was supported by The National Institutes of Health Grants R01-GM057391 (JS) and R01-GM098894 (QZ and JS), the Natural Science and Engineering Research Council of Canada (JEB, Discovery grant NSERC-2020-04241) and the Michael Smith Foundation for Health Research (JEB, Scholar Award 17686). ES-P was supported by a National Science Foundation Graduate Research Fellowship under Grant No. DGE-1650116.

The content is solely the responsibility of the authors and does not necessarily represent the official views of the National Institutes of Health.

---

**Table S1.**

| 693 |  |  |  |  |
| --- | --- | --- | --- | --- |
| **Data Set** | **PLCγ-1(CD)** | **PLCγ-1(CD) + FGFR** | **PLCγ-1(CD) + Lipid** | **PLCγ-1(CD) + FGFR + Lipid** |
| HDX reaction details | %D<sub>2</sub>O=72%<br>pH<sub>(read)</sub>= 7.0<br>Temp= 18°C | %D<sub>2</sub>O=72%<br>pH<sub>(read)</sub>= 7.0<br>Temp= 18°C | %D<sub>2</sub>O=72%<br>pH<sub>(read)</sub>= 7.0<br>Temp= 18°C | %D<sub>2</sub>O=72%<br>pH<sub>(read)</sub>= 7.0<br>Temp= 18°C |
| HDX time course | 3s, 30s, 300s, 3000s, 10000s | 3s, 30s, 300s, 3000s, 10000s | 3s, 30s, 300s, 3000s, 10000s | 3s, 30s, 300s, 3000s, 10000s |
| HDX controls | N/A | N/A | N/A | N/A |
| Back-exchange | Corrected based on %D<sub>2</sub>O | Corrected based on %D<sub>2</sub>O | Corrected based on %D<sub>2</sub>O | Corrected based on %D<sub>2</sub>O |
| Number of peptides | 254 | 254 | 254 | 254 |
| Sequence coverage | 91.5% | 91.5% | 91.5% | 91.5% |
| Average peptide length/ redundancy | Length = 13.0<br>Redundancy = 2.7 | Length = 13.0<br>Redundancy = 2.7 | Length = 13.0<br>Redundancy = 2.7 | Length = 13.0<br>Redundancy = 2.7 |
| Replicates | 3 | 3 | 3 | 3 |
| Repeatability | Average StDev = 0.4% | Average StDev = 0.4% | Average StDev = 0.5% | Average StDev = 0.5% |
| Significant differences in HDX | >5% and >0.4 Da and unpaired t-test <0.01 | >5% and >0.4 Da and unpaired t-test <0.01 | >5% and >0.4 Da and unpaired t-test <0.01 | >5% and >0.4 Da and unpaired t-test <0.01 |
| **Data Set** | **PLCγ-1(D1165)** | **PLCγ-1(D1165) + FGFR** | **PLCγ-1(D1165) + Lipid** | **PLCγ-1(D1165) + FGFR + Lipid** |
| HDX reaction details | %D<sub>2</sub>O=72%<br>pH<sub>(read)</sub>= 7.0<br>Temp= 18°C | %D<sub>2</sub>O=72%<br>pH<sub>(read)</sub>= 7.0<br>Temp= 18°C | %D<sub>2</sub>O=72%<br>pH<sub>(read)</sub>= 7.0<br>Temp= 18°C | %D<sub>2</sub>O=72%<br>pH<sub>(read)</sub>= 7.0<br>Temp= 18°C |
| HDX time course | 3s, 30s, 300s, 3000s, 10000s | 3s, 30s, 300s, 3000s, 10000s | 3s, 30s, 300s, 3000s, 10000s | 3s, 30s, 300s, 3000s, 10000s |
| HDX controls | N/A | N/A | N/A | N/A |
| Back-exchange | Corrected based on %D<sub>2</sub>O | Corrected based on %D<sub>2</sub>O | Corrected based on %D<sub>2</sub>O | Corrected based on %D<sub>2</sub>O |
| Number of peptides | 254 | 254 | 254 | 254 |
| Sequence coverage | 91.5% | 91.5% | 91.5% | 91.5% |
| Average peptide length/ redundancy | Length = 13.0<br>Redundancy = 2.7 | Length = 13.0<br>Redundancy = 2.7 | Length = 13.0<br>Redundancy = 2.7 | Length = 13.0<br>Redundancy = 2.7 |
| Replicates | 3 | 3 | 3 | 3 |
| Repeatability | Average StDev = 0.4% | Average StDev = 0.4% | Average StDev = 0.4% | Average StDev = 0.4% |
| Significant differences in HDX | >5% and >0.4 Da and unpaired t-test <0.01 | >5% and >0.4 Da and unpaired t-test <0.01 | >5% and >0.4 Da and unpaired t-test <0.01 | >5% and >0.4 Da and unpaired t-test <0.01 |

**Figure S1.**
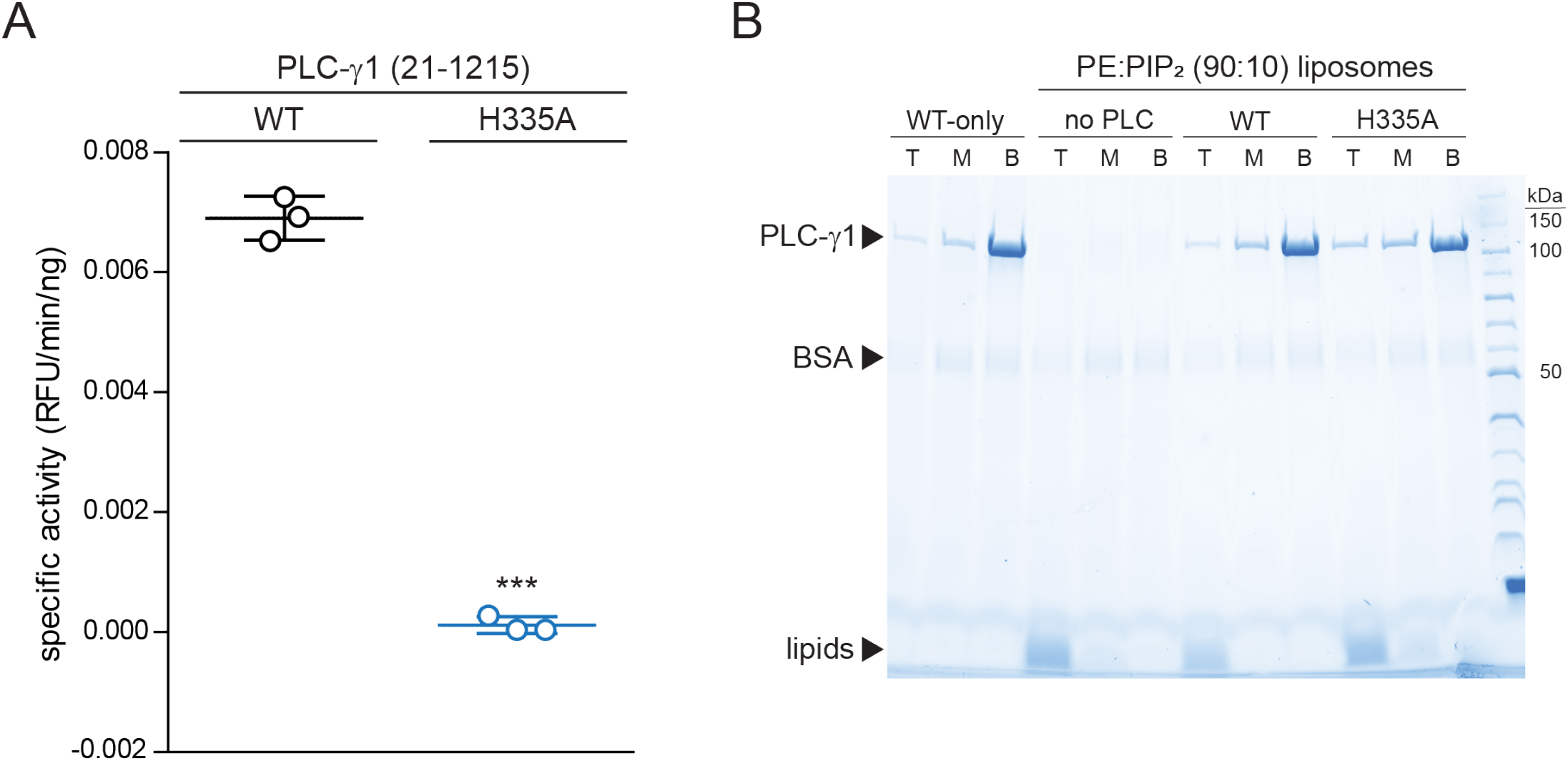
PLC-γ1 (H335A) is catalytically inactive while retaining capacity to binds liposomes. (**A**) Specific activities measured with the membrane-embedded substrate XY-69. XY-69 (0.5μM) was incorporated into liposomes containing PE:PIP_2_ (80:20) prior to addition of 10 nM of wild-type (WT) PLC-γ1 or PLC-γ1 (H335A). Phospholipase activities were determined by quantifying XY-69 hydrolysis in real-time and presented as the mean ± SD of three independent experiments, each with three or more technical replicates. Statistical significance was determined with a t-test represented as *** for a p < 0.05. (**B**) Membrane floatation assay. Indicated proteins were incubated with liposomes of PE:PIP_2_:NBD-PE (89.8:10:0.2) and centrifuged in a sucrose gradient (0-30%). After centrifugation, top (T), middle (M), and bottom (B) fractions were collected followed by SDS-PAGE and staining with Coomassie Brilliant Blue. (**C**) Deuterium incorporation was measured for PLC-γ1 (H335A) and differences in deuterium incorporation were calculated relative to wild-type PLC-γ1. Only peptides encompassing (H335A) showed significant differences (>5% and >0.4 Da difference in exchange at any time point using an unpaired Student’s t-test value p<0.01) and this region was mapped onto the structure. Percent deuterium incorporation for this region is graphed versus time (3, 30, 300, 3000 sec) and shown next to the structure as the mean ± SD (n = 3); most standard deviations are smaller than the size of the corresponding point. The highest percentage of deuterium incorporation is shown above the curve and an asterisk (*) above a time point indicates a difference >0.4 Da for that time point.

**Figure S2.**
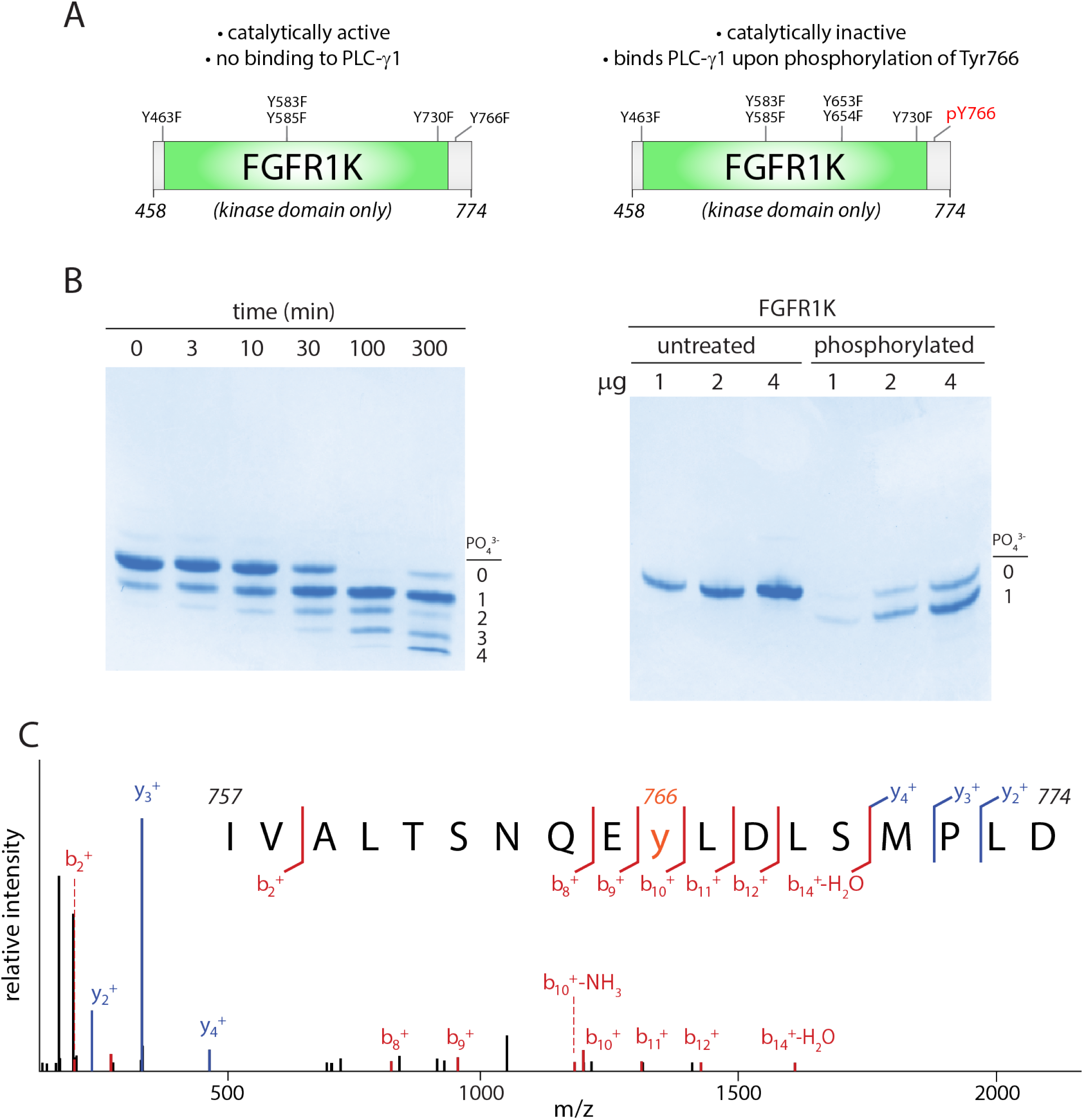
FGFR1K specifically phosphorylated on Tyr766. (**A**) Schematic diagram of the kinase domain (residues 458-774) of FGFR1K. The six tyrosines mutated to phenylalanine to prevent phosphorylation and render the kinase resistant to phosphorylation-dependent activation are listed. Phosphorylation of Tyr766 is shown in red. (**B**) Time course of phosphorylation (left) and preparative phosphorylation (right) of catalytically-inactive FGFR1K (100 μM) using equimolar concentrations of an equivalent, catalytically-active fragment. Reactions were terminated by separating the two forms of FGFR1K using affinity chromatography; native PAGE was subsequently used to separate phosphorylated forms of the catalytically-inactive version followed by staining with Coomassie Brilliant Blue. Preparative phosphorylation was carried out for 100 min. (**C**) Catalytically-inactive FGFR1K was phosphorylated specifically at Tyr766 as determined by LC-MS/MS.

**Figure S3.**
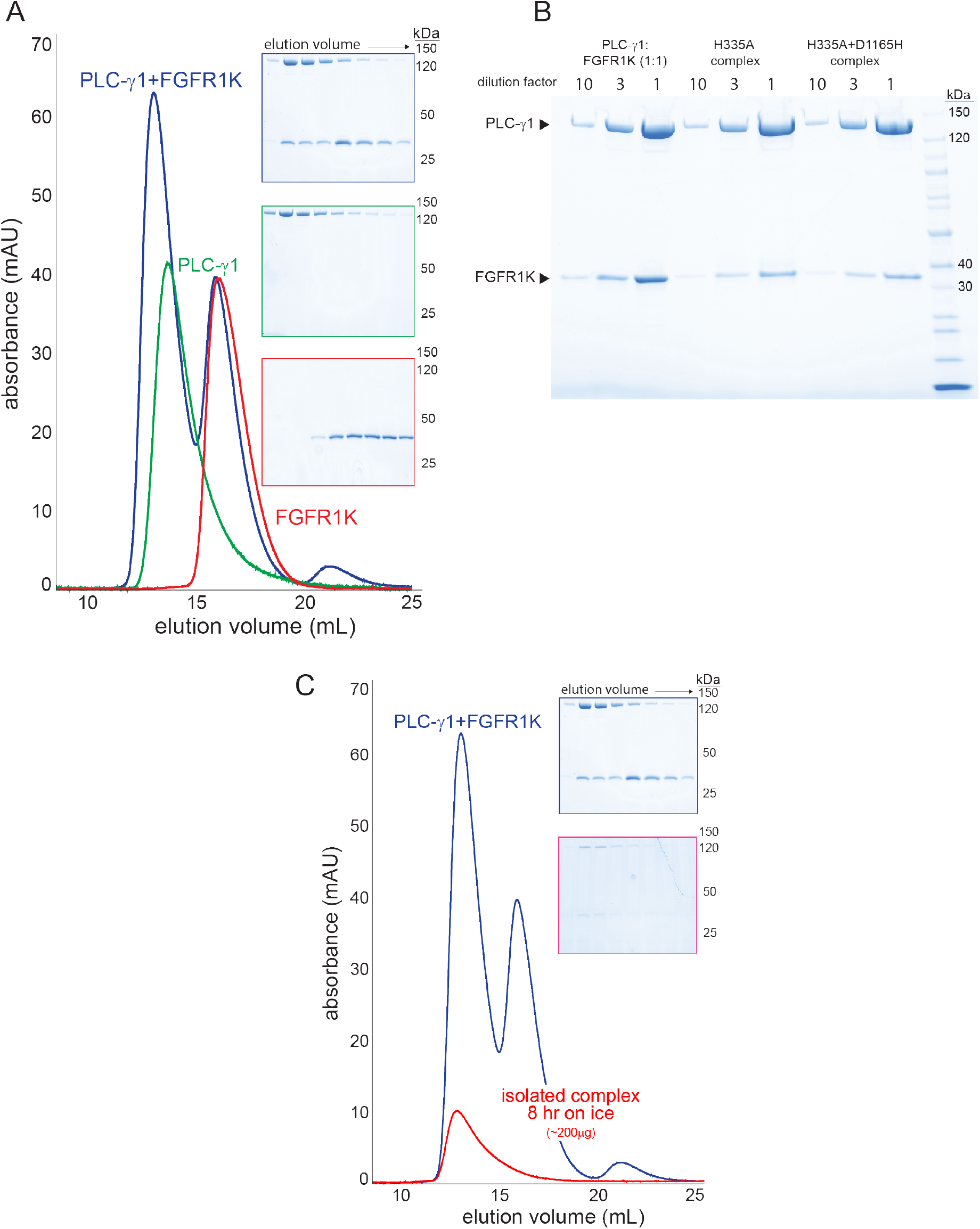
Stable complexes of PLC-γ1 and phosphorylated FGFR1K. (**A**) PLC-γ1 (H335A) or PLC-γ1 (H335A+D1165H) were mixed with a 2-fold molar excess of phosphorylated FGFR1K and isolated by size exclusion chromatography; representative chromatograms are shown with fractions visualized after SDS-PAGE and staining with Coomassie Brilliant Blue (insets). (**B**) Final complexes were visualized similarly. For reference, a 1:1 mixture of PLC-γ1 and FGFR1K was also visualized. (**C**) Stability of the complex of PLC-γ1 (H335A) and FGFR1K was assessed by size exclusion chromatography after 8 hours on ice (red curve). Blue curve is duplicated from panel A and shown for reference.

**Figure S4.**
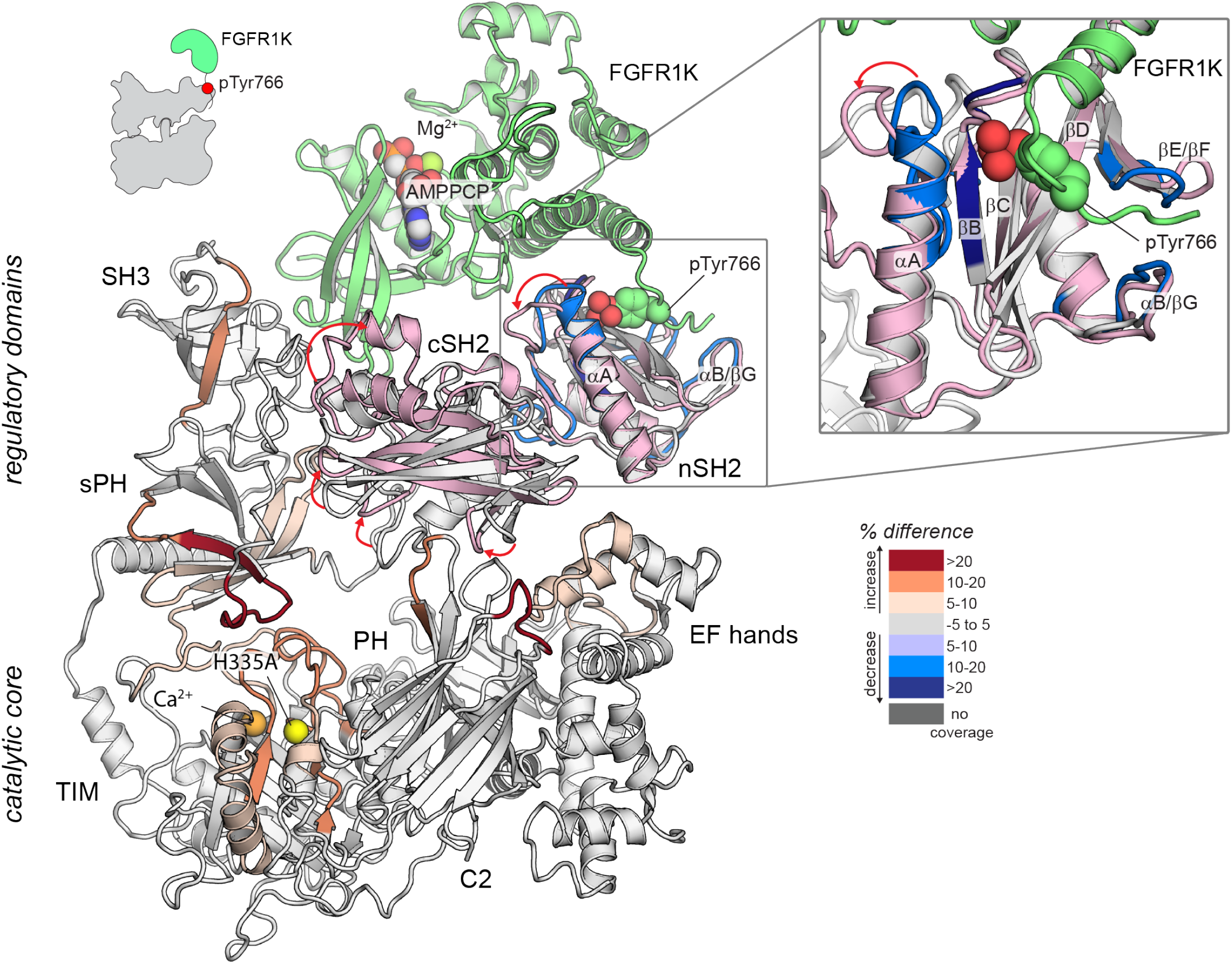
Potential mechanism of priming of PLC-γ1 by FGFR1K. Structural superimposition of autoinhibited PLC-γ1 (PDB code: 6PBC) and the tandem SH2 domains of PLC-γ1 (light pink) bound to FGFR1K (green; PDB code: 3GQI). Structures superimposed using the nSH2 domains only. Autoinhibited PLC-γ1 is colored gray with regions of differential exchange upon binding FGFR1K colored as in Figure 2A. AMPPCP is a nonhydrolyzable analog of ATP. Red arrows highlight changes in the position of loops within the tandem SH2 domains.

**Figure S5.**
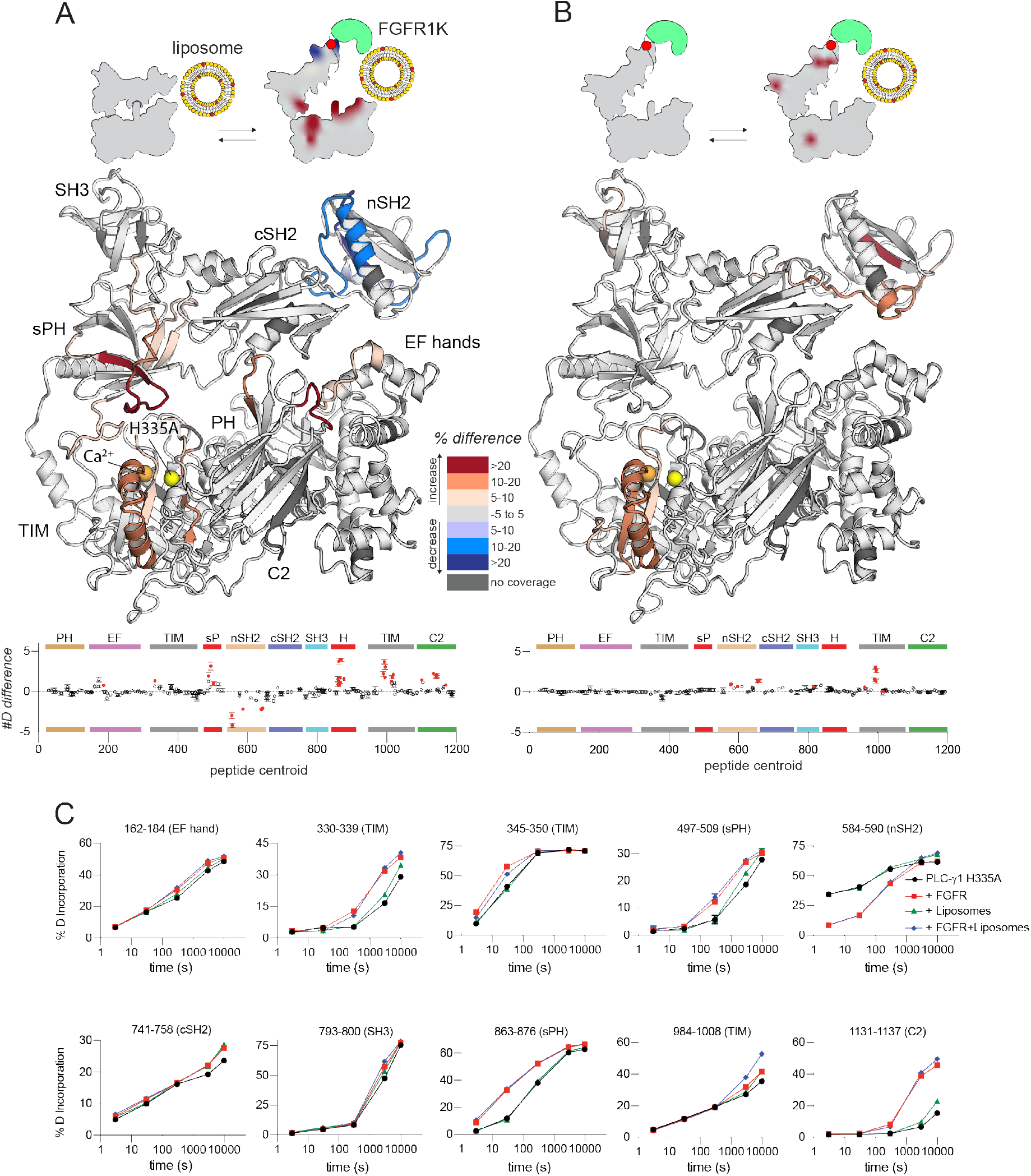
FGFR1K and liposomes act essentially independently to affect sites of exchange within PLC-γ1. Differences in deuterium incorporation was measured for PLC-γ1 (H335A) bound to FGFR1K and in the presence of liposomes containing PE:PIP_2_ (90:10). Differences in deuterium incorporation were calculated relative to PLC-γ1 (H335A) in the presence of PE:PIP_2_ (90:10) liposomes (**A**) or PLC-γ1 (H335A) bound to FGFR1K (**B**). Peptides that showed significant difference in deuterium incorporation (>5% and >0.4 Da difference in exchange at any time point, with an unpaired Student’s t-test value <0.01) are mapped onto the structure. For each comparison, the difference in the number of incorporated deuterons was averaged over all time points and shown below the structures as the mean ± SD (n = 3). Each circle represents the central residue of a corresponding peptide; red circles indicate peptides with a significant difference between conditions. (**C**) Percent deuterium incorporation for select peptides spanning PLC-γ1 (H335A). Percent deuterium incorporation is shown for PLC-γ1 (H335A) alone (Apo; black line), bound to FGFR1K (red line), in the presence of PE:PIP_2_ (90:10) liposomes (green line), and with both kinase and liposomes (blue line). Percent deuterium incorporation is the mean ± SD (n=3), with most SD smaller than the size of the point. The full deuterium incorporation data for all peptides are shown in the source data.

**Figure S6.**
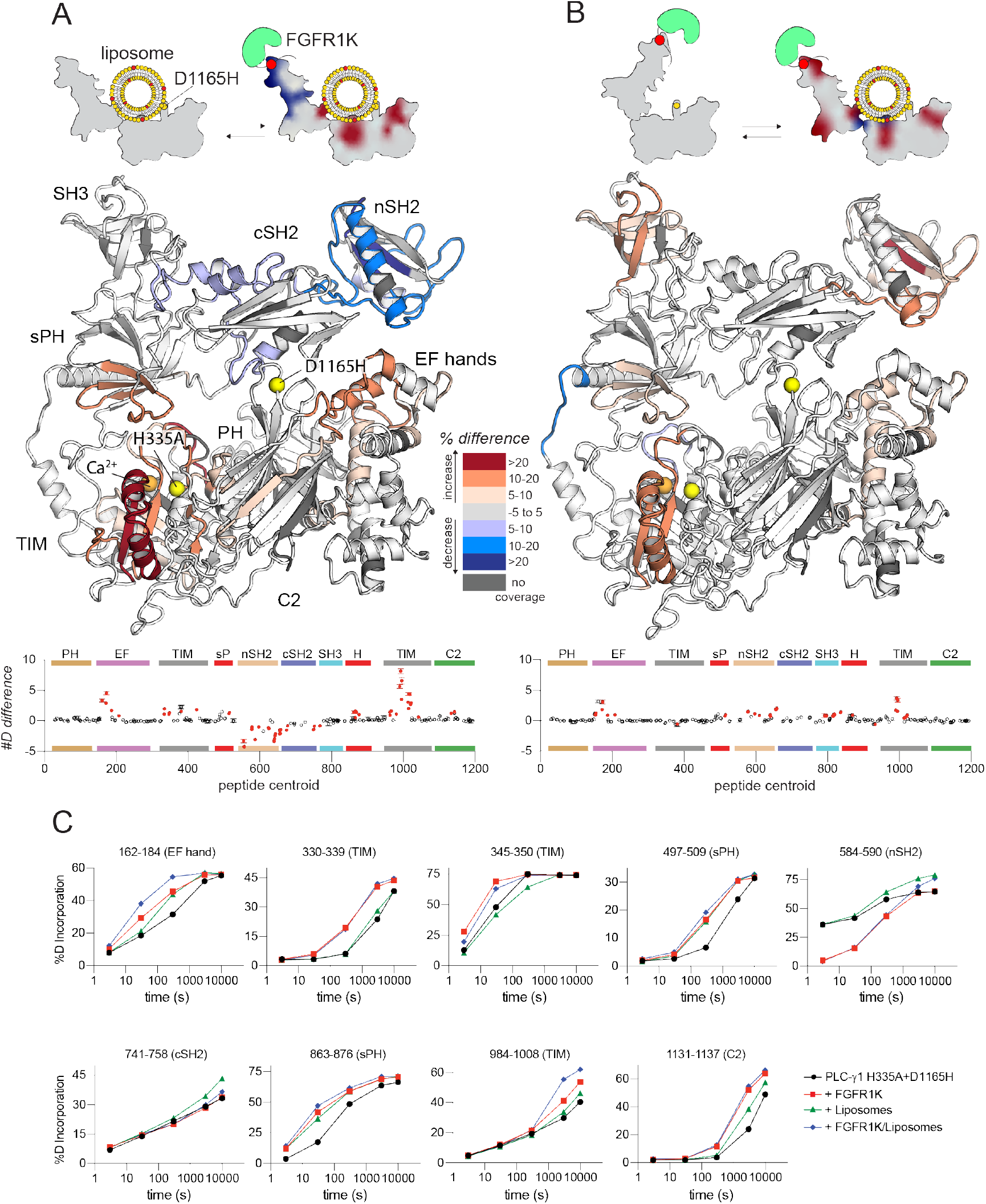
FGFR1K and liposomes cooperate to affect the deuterium exchange of PLC-γ1 (D1165H). Deuterium incorporation was measured for PLC-γ1 (H335A+D1165H) bound to FGFR1K in the presence of PE:PIP_2_ (90:10) liposomes. Differences in deuterium incorporation were calculated relative to PLC-γ1 (H335A+D1165H) in the presence of PE:PIP_2_ liposomes (**A**) or bound to FGFR1K (**B**). Peptides that showed significant H/D exchange differences (by the following criteria: >5% and >0.4 Da difference in exchange at any time point (3, 30, 300, 3000, and 10000 s), with an unpaired Student’s t-test value <0.01) are mapped onto the structure. For each comparison, the difference in the number of incorporated deuterons was averaged over all time points and shown below the structures as the mean ± SD (n = 3). Each circle represents the central residue of a corresponding peptide; red circles indicate peptides with a significant difference between conditions. (**C**) Percent deuterium incorporation for select peptides of PLC-γ1 (H335A+D1165H) plotted as the mean ± SD (n=3) for the indicated conditions. Most error bars are smaller than the size of the corresponding point. The full deuterium incorporation data for all peptides are shown in the source data.

**Figure S7.**
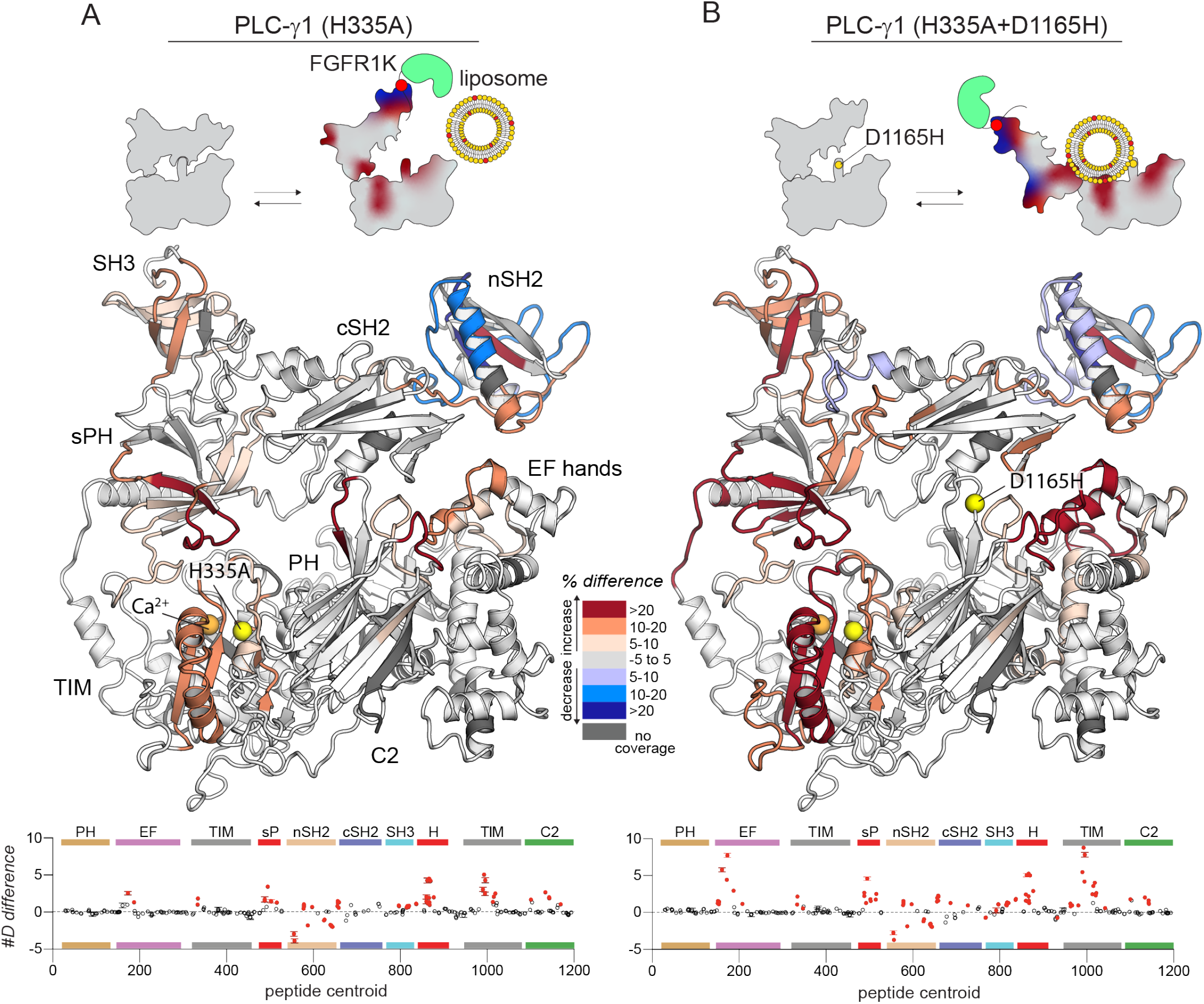
FGFR1K and liposomes have enhanced capacity to affect the deuterium exchange of the oncogenic mutant of PLC-γ1 compared to the WT. Differences in deuterium incorporation was measured for PLC-γ1 (H335A) or PLC-γ1 (H335A+D1165H) with both kinase and liposomes. Differences in deuterium incorporation were calculated relative to PLC-γ1 (H335A) alone (**A**) or to PLC-γ1 (H335A+D1165H) alone (**B**). Peptides that showed significant H/D exchange differences (by the following criteria: >5% and >0.4 Da difference in exchange at any time point (3, 30, 300, 3000, and 10000 s), with an unpaired Student’s t-test value <0.01) are mapped onto the structure. For each comparison, the difference in the number of incorporated deuterons was averaged over all time points and shown below the structures as the mean ± SD (n = 3). Each circle represents the central residue of a corresponding peptide; red circles indicate peptides with a significant difference between conditions. Figure duplicates portions of figures 2 and 4 to provide a direct comparison of the effects of FGFR1 and liposomes on the exchange profiles of PLC-γ1 (H335A) and (H335A + D1165H).

## References

Arteaga, C.L., Johnson, M.D., Todderud, G., Coffey, R.J., Carpenter, G., and Page, D.L. (1991). Elevated content of the tyrosine kinase substrate phospholipase C-γ1 in primary human breast carcinomas. Proc Natl Acad Sci USA 88, 10435–10439.

Asokan, S.B., Johnson, H.E., Rahman, A., King, S.J., Rotty, J.D., Lebedeva, I.P., Haugh, J.M., and Bear, J.E. (2014). Mesenchymal chemotaxis requires selective inactivation of myosin II at the leading edge via a noncanonical PLCγ/PKCα pathway. Dev Cell 31, 747–760.

Bae, J.H., Lew, E.D., Yuzawa, S., Tome, F., Lax, I., and Schlessinger, J. (2009). The selectivity of receptor tyrosine kinase signaling is controlled by a secondary SH2 domain binding site. Cell 138, 514–524.

Bunney, T.D., Esposito, D., Mas-Droux, C., Lamber, E., Baxendale, R.W., Martins, M., Cole, A., Svergun, D., Driscoll, P.C., and Katan, M. (2012). Structural and functional integration of the PLCγ interaction domains critical for regulatory mechanisms and signaling deregulation. Structure 20, 2062–2075.

Bunney, T.D., Opaleye, O., Roe, S.M., Vatter, P., Baxendale, R.W., Walliser, C., Everett, K.L., Josephs, M.B., Christow, C., Rodrigues-Lima, F., et al. (2009). Structural insights into formation of an active signaling complex between Rac and phospholipase Cγ2. Mol Cell 34, 223–233.

Burgess, W.H., Dionne, C.A., Kaplow, J., Mudd, R., Friesel, R., Zilberstein, A., Schlessinger, J., and Jaye, M. (1990). Characterization and cDNA cloning of phospholipase C-γ, a major substrate for heparin-binding growth factor 1 (acidic fibroblast growth factor)-activated tyrosine kinase. Mol Cell Biol 10, 4770–4777.

Chattopadhyay, A., Vecchi, M., Ji, Q., Mernaugh, R., and Carpenter, G. (1999). The role of individual SH2 domains in mediating association of phospholipase C-γ1 with the activated EGF receptor. J Biol Chem 274, 26091–26097.

DeBell, K., Graham, L., Reischl, I., Serrano, C., Bonvini, E., and Rellahan, B. (2007). Intramolecular regulation of phospholipase C-γ1 by its C-terminal Src homology 2 domain. Mol Cell Biol 27, 854–863.

DeBell, K.E., Stoica, B.A., Veri, M.C., Di Baldassarre, A., Miscia, S., Graham, L.J., Rellahan, B.L., Ishiai, M., Kurosaki, T., and Bonvini, E. (1999). Functional independence and interdependence of the Src homology domains of phospholipase C-γ1 in B-cell receptor signal transduction. Mol Cell Biol 19, 7388–7398.

Dobbs, J.M., Jenkins, M.L., and Burke, J.E. (2020). Escherichia coli and Sf9 contaminant databases to increase efficiency of tandem mass spectrometry peptide identification in structural mass spectrometry experiments. J Am Soc Mass Spectrom 31, 2202–2209.

Ellis, M.V., James, S.R., Perisic, O., Downes, C.P., Williams, R.L., and Katan, M. (1998). Catalytic domain of phosphoinositide-specific phospholipase C (PLC). Mutational analysis of residues within the active site and hydrophobic ridge of PLCδ1. J Biol Chem 273, 11650–11659.

Eswarakumar, V.P., Lax, I., and Schlessinger, J. (2005). Cellular signaling by fibroblast growth factor receptors. Cytokine Growth Factor Rev 16, 139–149.

Gresset, A., Hicks, S.N., Harden, T.K., and Sondek, J. (2010). Mechanism of phosphorylation-induced activation of phospholipase C-γ isozymes. J Biol Chem 285, 35836–35847.

Gresset, A., Sondek, J., and Harden, T.K. (2012). The phospholipase C isozymes and their regulation. Subcell Biochem 58, 61–94.

Hajicek, N., Charpentier, T.H., Rush, J.R., Harden, T.K., and Sondek, J. (2013). Autoinhibition and phosphorylation-induced activation of phospholipase C-γ isozymes. Biochem 52, 4810–4819.

Hajicek, N., Keith, N.C., Siraliev-Perez, E., Temple, B.R., Huang, W., Zhang, Q., Harden, T.K., and Sondek, J. (2019). Structural basis for the activation of PLC-γ isozymes by phosphorylation and cancer-associated mutations. eLife 8, e51700.

Horstman, D.A., Chattopadhyay, A., and Carpenter, G. (1999). The influence of deletion mutations on phospholipase C-γ1 activity. Arch Biochem Biophys 361, 149–155.

Huang, W., Hicks, S.N., Sondek, J., and Zhang, Q. (2011). A fluorogenic, small molecule reporter for mammalian phospholipase C isozymes. ACS Chem Biol 6, 223–228.

Huang, W., Wang, X., Endo-Streeter, S., Barrett, M., Waybright, J., Wohlfeld, C., Hajicek, N., Harden, T.K., Sondek, J., and Zhang, Q. (2018). A membrane-associated, fluorogenic reporter for mammalian phospholipase C isozymes. J Biol Chem 293, 1728–1735.

Ji, Q.S., Chattopadhyay, A., Vecchi, M., and Carpenter, G. (1999). Physiological requirement for both SH2 domains for phospholipase C-γ1 function and interaction with platelet-derived growth factor receptors. Mol Cell Biol 19, 4961–4970.

Kadamur, G., and Ross, E.M. (2013). Mammalian phospholipase C. Annu Rev Physiol 75, 127–154.

Kataoka, K., Nagata, Y., Kitanaka, A., Shiraishi, Y., Shimamura, T., Yasunaga, J., Totoki, Y., Chiba, K., Sato-Otsubo, A., Nagae, G., et al. (2015). Integrated molecular analysis of adult T cell leukemia/lymphoma. Nat Genet 47, 1304–1315.

Kim, H.K., Kim, J.W., Zilberstein, A., Margolis, B., Kim, J.G., Schlessinger, J., and Rhee, S.G. (1991). PDGF stimulation of inositol phospholipid hydrolysis requires PLC-γ1 phosphorylation on tyrosine residues 783 and 1254. Cell 65, 435–441.

Kim, J.W., Sim, S.S., Kim, U.H., Nishibe, S., Wahl, M.I., Carpenter, G., and Rhee, S.G. (1990). Tyrosine residues in bovine phospholipase C-γ phosphorylated by the epidermal growth factor receptor in vitro. J Biol Chem 265, 3940–3943.

Kim, M.J., Chang, J.S., Park, S.K., Hwang, J.I., Ryu, S.H., and Suh, P.G. (2000). Direct interaction of SOS1 Ras exchange protein with the SH3 domain of phospholipase C-γ1. Biochem 39, 8674–8682.

Kleineidam, L., Chouraki, V., Prochnicki, T., van der Lee, S.J., Madrid-Marquez, L., Wagner-Thelen, H., Karaca, I., Weinhold, L., Wolfsgruber, S., Boland, A., et al. (2020). PLCG2 protective variant p.P522R modulates tau pathology and disease progression in patients with mild cognitive impairment. Acta Neuropathol 139, 1025–1044.

Koss, H., Bunney, T.D., Behjati, S., and Katan, M. (2014). Dysfunction of phospholipase Cγ in immune disorders and cancer. Trends Biochem Sci 39, 603–611.

Law, C.L., Chandran, K.A., Sidorenko, S.P., and Clark, E.A. (1996). Phospholipase C-γ1 interacts with conserved phosphotyrosyl residues in the linker region of Syk and is a substrate for Syk. Mol Cell Biol 16, 1305–1315.

Liu, Y., Bunney, T.D., Khosa, S., Mace, K., Beckenbauer, K., Askwith, T., Maslen, S., Stubbs, C., de Oliveira, T.M., Sader, K., et al. (2020). Structural insights and activating mutations in diverse pathologies define mechanisms of deregulation for phospholipase C γ enzymes. EBioMedicine 51, 102607.

Manna, A., Zhao, H., Wada, J., Balagopalan, L., Tagad, H.D., Appella, E., Schuck, P., and Samelson, L.E. (2018). Cooperative assembly of a four-molecule signaling complex formed upon T cell antigen receptor activation. Proc Natl Acad Sci USA 115, E11914–E11923.

Masson, G.R., Burke, J.E., Ahn, N.G., Anand, G.S., Borchers, C., Brier, S., Bou-Assaf, G.M., Engen, J.R., Englander, S.W., Faber, J., et al. (2019). Recommendations for performing, interpreting and reporting hydrogen deuterium exchange mass spectrometry (HDX-MS) experiments. Nat Methods 16, 595–602.

Mohammadi, M., Honegger, A.M., Rotin, D., Fischer, R., Bellot, F., Li, W., Dionne, C.A., Jaye, M., Rubinstein, M., and Schlessinger, J. (1991). A tyrosine-phosphorylated carboxy-terminal peptide of the fibroblast growth factor receptor (Flg) is a binding site for the SH2 domain of phospholipase C-γ1. Mol Cell Biol 11, 5068–5078.

Nakanishi, O., Shibasaki, F., Hidaka, M., Homma, Y., and Takenawa, T. (1993). Phospholipase C-γ1 associates with viral and cellular src kinases. J Biol Chem 268, 10754–10759.

Nishibe, S., Wahl, M.I., Hernandez-Sotomayor, S.M., Tonks, N.K., Rhee, S.G., and Carpenter, G. (1990). Increase of the catalytic activity of phospholipase C-γ1 by tyrosine phosphorylation. Science 250, 1253–1256.

Noh, D.Y., Shin, S.H., and Rhee, S.G. (1995). Phosphoinositide-specific phospholipase C and mitogenic signaling. Biochim Biophys Acta 1242, 99–113.

Patel, V.M., Flanagan, C.E., Martins, M., Jones, C.L., Butler, R.M., Woollard, W.J., Bakr, F.S., Yoxall, A., Begum, N., Katan, M., et al. (2020). Frequent and persistent PLCG1 mutations in Sezary cells directly enhance PLCγ1 activity and stimulate NFκB, AP-1, and NFAT signaling. J Invest Dermatol 140, 380–389 e384.

Perez-Riverol, Y., Csordas, A., Bai, J., Bernal-Llinares, M., Hewapathirana, S., Kundu, D.J., Inuganti, A., Griss, J., Mayer, G., Eisenacher, M., et al. (2019). The PRIDE database and related tools and resources in 2019: improving support for quantification data. Nucleic Acids Res 47, D442–D450.

Piechulek, T., Rehlen, T., Walliser, C., Vatter, P., Moepps, B., and Gierschik, P. (2005). Isozyme-specific stimulation of phospholipase C-γ2 by Rac GTPases. J Biol Chem 280, 38923–38931.

Poulin, B., Sekiya, F., and Rhee, S.G. (2000). Differential roles of the Src homology 2 domains of phospholipase C-γ1 (PLC-γ1) in platelet-derived growth factor-induced activation of PLC-γ1 in intact cells. J Biol Chem 275, 6411–6416.

Poulin, B., Sekiya, F., and Rhee, S.G. (2005). Intramolecular interaction between phosphorylated tyrosine-783 and the C-terminal Src homology 2 domain activates phospholipase C-γ1. Proc Natl Acad Sci USA 102, 4276–4281.

Rouquette-Jazdanian, A.K., Sommers, C.L., Kortum, R.L., Morrison, D.K., and Samelson, L.E. (2012). LAT-independent Erk activation via Bam32-PLC-γ1-Pak1 complexes: GTPase-independent Pak1 activation. Mol Cell 48, 298–312.

Schade, A., Walliser, C., Wist, M., Haas, J., Vatter, P., Kraus, J.M., Filingeri, D., Havenith, G., Kestler, H.A., Milner, J.D., et al. (2016). Cool-temperature-mediated activation of phospholipase C-γ2 in the human hereditary disease PLAID. Cell Signal 28, 1237–1251.

Schaeffer, E.M., Debnath, J., Yap, G., McVicar, D., Liao, X.C., Littman, D.R., Sher, A., Varmus, H.E., Lenardo, M.J., and Schwartzberg, P.L. (1999). Requirement for Tec kinases Rlk and Itk in T cell receptor signaling and immunity. Science 284, 638–641.

Sims, R., van der Lee, S.J., Naj, A.C., Bellenguez, C., Badarinarayan, N., Jakobsdottir, J., Kunkle, B.W., Boland, A., Raybould, R., Bis, J.C., et al. (2017). Rare coding variants in PLCG2, ABI3, and TREM2 implicate microglial-mediated innate immunity in Alzheimer’s disease. Nat Genet 49, 1373–1384.

Stariha, J.T.B., Hoffmann, R.M., Hamelin, D.J., and Burke, J.E. (2021). Probing protein-membrane interactions and dynamics using hydrogen-deuterium exchange mass spectrometry (HDX-MS). Methods Mol Biol 2263, 465–485.

Tvorogov, D., and Carpenter, G. (2002). EGF-dependent association of phospholipase C-γ1 with c-Cbl. Exp Cell Res 277, 86–94.

Vetter, M.L., Martin-Zanca, D., Parada, L.F., Bishop, J.M., and Kaplan, D.R. (1991). Nerve growth factor rapidly stimulates tyrosine phosphorylation of phospholipase C-γ1 by a kinase activity associated with the product of the trk protooncogene. Proc Natl Acad Sci USA 88, 5650–5654.

Walliser, C., Retlich, M., Harris, R., Everett, K.L., Josephs, M.B., Vatter, P., Esposito, D., Driscoll, P.C., Katan, M., Gierschik, P., et al. (2008). Rac regulates its effector phospholipase Cγ2 through interaction with a split pleckstrin homology domain. J Biol Chem 283, 30351–30362.

Watanabe, D., Hashimoto, S., Ishiai, M., Matsushita, M., Baba, Y., Kishimoto, T., Kurosaki, T., and Tsukada, S. (2001). Four tyrosine residues in phospholipase C-γ2, identified as Btk-dependent phosphorylation sites, are required for B cell antigen receptor-coupled calcium signaling. J Biol Chem 276, 38595–38601.

Woyach, J.A., Furman, R.R., Liu, T.M., Ozer, H.G., Zapatka, M., Ruppert, A.S., Xue, L., Li, D.H., Steggerda, S.M., Versele, M., et al. (2014). Resistance mechanisms for the Bruton’s tyrosine kinase inhibitor ibrutinib. New Engl J Med 370, 2286–2294.

Yablonski, D., Kadlecek, T., and Weiss, A. (2001). Identification of a phospholipase C-γ1 (PLC-γ1) SH3 domain-binding site in SLP-76 required for T-cell receptor-mediated activation of PLC-γ1 and NFAT. Mol Cell Biol 21, 4208–4218.

